# Firing rate adaptation in grid cells produces macroscopic hexadirectional modulation

**DOI:** 10.64898/2026.10.19.760191

**Authors:** Guglielmo Reggio, Misun Kim, Zilong Ji, Daniel Bush, Caswell Barry, Neil Burgess

**Affiliations:** UCL Institute of Cognitive Neuroscience, London, UK; UCL Queen Square Institute of Neurology, London, UK; Department of Neuroscience, Physiology and Pharmacology, University College London, UK; Research Department of Cell and Developmental Biology, University College London, UK

**Author notes:** These authors contributed equally to this work (Guglielmo Reggio, Misun Kim, Neil Burgess).

## Abstract

Hexadirectional modulation (HexMod) of macroscopic neural activity by movement direction is widely used to measure grid cell activity in human neuroimaging, but its neuronal origin is unclear. Two mechanisms have been proposed: conjunctive grid×direction cells whose preferred directions align with the grid axes, and firing rate adaptation, which reduces firing when grid fields are visited in quick succession during movement along grid axes. Analysing almost 4,000 grid cells from foraging rats, we found that conjunctive cells’ directional tuning was broad and unaligned with grid axes, whereas firing was reduced shortly after grid field visits, consistent with firing rate adaptation. Simulations of realistic grid cell populations demonstrate HexMod with the typical trajectories of human experiments, and show how it depends on grid scale, adaptation time constant, movement speed, and controlling for variation with location. We show firing rate adaptation provides a plausible mechanism for HexMod and a framework for relating human neuroimaging to neuronal activity.

## Introduction

Grid cells in the medial entorhinal cortex (MEC) exhibit spatially periodic firing fields that form a hexagonal lattice across the environment, providing a metric for path integration and spatial memory (Hafting et al., 2005; McNaughton et al., 2006). Since their discovery, it has been proposed that grid cells support diverse human cognitive functions beyond spatial navigation such as imagination, inference, and conceptual organisation; see review (Behrens et al., 2018; Bellmund et al., 2018).

It has been proposed that grid cell activity in humans and non-human primates can be measured non-invasively by looking for the modulation of macroscopic neural activity by running direction with six-fold rotation symmetry (Doeller et al., 2010). This method has subsequently been applied across a range of tasks from multiple laboratories, e.g. (Bao et al., 2019; Bongioanni et al., 2021; Constantinescu et al., 2016; Park et al., 2021; Qu et al., 2026; Stangl et al., 2018). This “hexadirectional modulation” (HexMod) relies on the assumption that populations of grid cells share a similar grid orientation, even if they differ in scale (Doeller et al., 2010; Stensola et al., 2012) (Fig 1A,B). Two mechanisms have been proposed to underlie HexMod and assessed theoretically (Bin Khalid et al., 2024; Doeller et al., 2010). First, conjunctive grid × head-direction cells (Sargolini et al., 2006) could fire more when an animal moves parallel to the grid axes, if their preferred firing directions cluster around these six directions and their directional tuning is narrow enough (Bin Khalid et al., 2024; Doeller et al., 2010) (Fig 1C). Second, grid cells could fire less when an animal’s movement direction aligns with the grid axes, if firing rate adaptation reduces activity when grid fields are visited in quick succession which can happen during straight aligned movements (Fig 1D).

**Figure 1.**
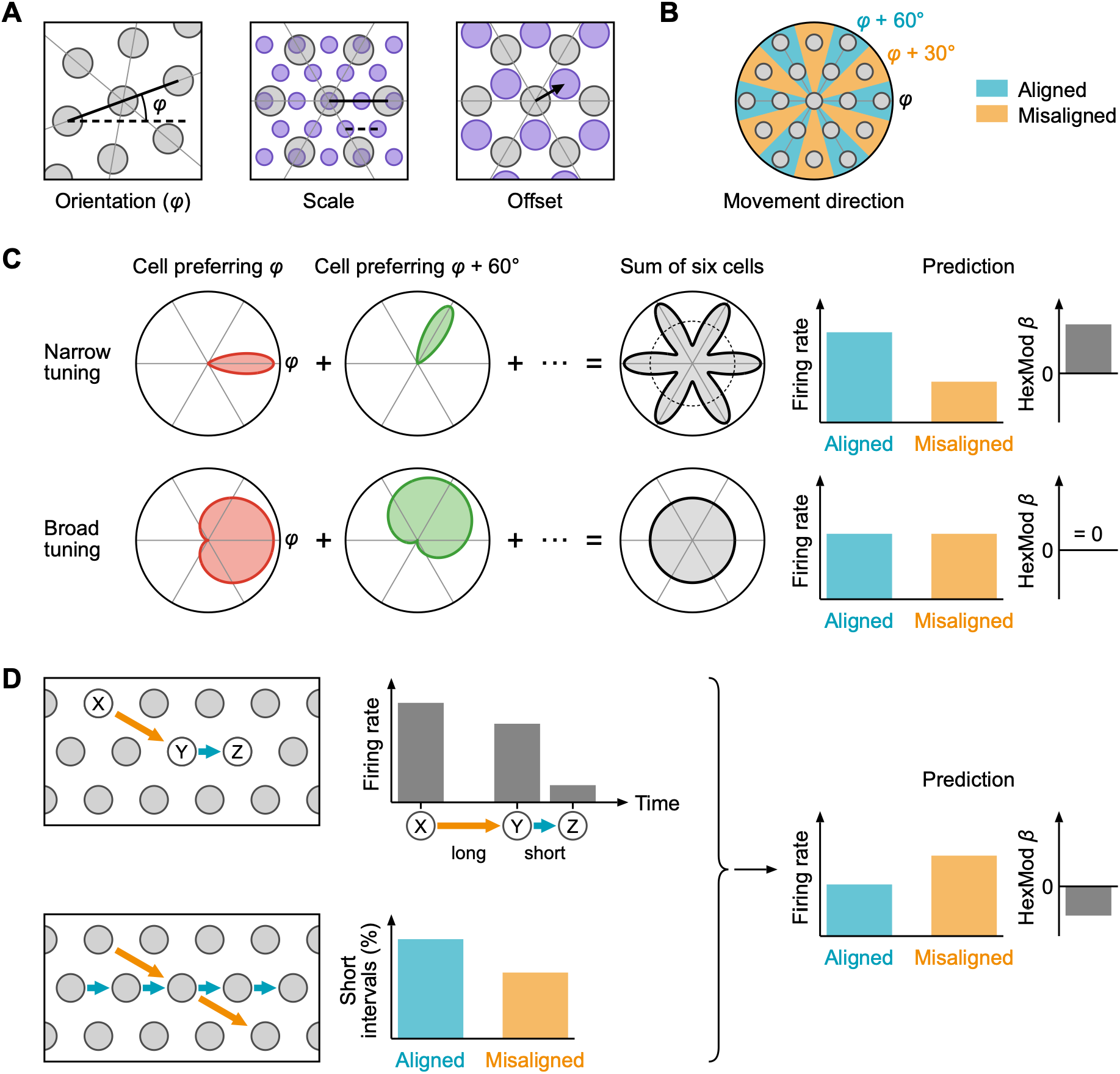
Potential neuronal mechanisms of hexadirectional modulation (HexMod). **A.** Grid cell properties. Grid cells of one module share the orientation (φ) and scale of their grid and differ in offset. Grey, one cell; purple, a second cell; thin lines, grid axes of the grey cell. **B.** HexMod: grid cell activity is modulated by whether the movement direction is aligned (blue; φ + k × 60°) or misaligned (orange; φ + 30° + k × 60°) with the grid axes. **C.** Conjunctive hypothesis. If the preferred firing directions of conjunctive grid × head-direction cells are aligned with the grid axes and their directional tuning is narrow (top row, von Mises function, standard deviation 15°), their summed activity is higher for aligned than misaligned movements, giving a positive regression coefficient β for a hexadirectional regressor (HexMod β). With broad tuning (bottom row, cosine tuning, SD= 67.5°), the sum of six cells with the same preferred directions is uniform and β is zero. Polar plots show firing rate against movement direction for two example cells (red, green) and for the sum of six such cells, one for each grid axis direction. Each cell’s tuning curve is scaled to its own peak, and the two sums are drawn on the same scale. **D.** Adaptation hypothesis predicts lower firing rates for aligned movement, and therefore a negative HexMod β. This arises because firing is more strongly suppressed after a short (Y → Z) than a long (X → Y) inter-field interval (top row), and aligned trajectories more often generate short inter-field intervals (bottom row), for straight movements. Circles are firing fields and arrows are movements from one field to the next (blue, aligned; orange, misaligned).

Distinguishing between the mechanisms underlying HexMod using empirical data is crucial for optimising experimental design. HexMod has been measured in large groups of participants to examine the maturity of grid cells during development (Qu et al., 2026), their decline during ageing (Stangl et al., 2018), and alterations in neurological and psychiatric conditions such as Alzheimer’s disease (Bierbrauer et al., 2020; Kunz et al., 2015) and schizophrenia (Convertino et al., 2023). Under the adaptation hypothesis, HexMod strength would be strongly affected by experimental parameters such as the tortuosity and length of trajectories (Bin Khalid et al., 2024), while interpreting grid orientation critically depends on the underlying mechanism, because the two hypotheses predict opposite signal modulations (either aligned or misaligned with the grid axes).

Here, we analysed almost 4,000 grid cells from 32 rats, drawn from six published datasets (see Methods) (Barry et al., 2012; Brun et al., 2008; Hafting et al., 2005; Ólafsdóttir et al., 2016; Sargolini et al., 2006; Vollan et al., 2025) while controlling for the effects of inhomogeneous behavioural sampling of location, running speed, and direction (Burgess et al., 2005). We show that conjunctive cells’ directional tuning is not aligned to grid axes, but that HexMod can arise from adaptation-driven suppression, depending on how movement trajectories sample grid firing patterns. We then simulate a population of realistic grid cells with the typical trajectories used in human neuroimaging experiments and show that HexMod strength is larger for smaller-scale grid firing patterns. Finally, we propose methods for enhancing the detection of HexMod by controlling for variations in activity with location and speed.

## Results

### Conjunctive grid × direction cells are broadly tuned and unaligned with grid axes

We first tested whether conjunctive cells had preferred directions aligned with the grid axes and directional tuning narrow enough to generate HexMod (Fig 1C). Of 1,171 conjunctive cells from 21 animals, 564 (48%) had a preferred direction within 15° of a grid axis, close to the 50% expected by chance (binomial test, p = 0.22; Fig 2A,B). Directional tuning was also broad: the median circular standard deviation was 73° (Fig 2C). At this width, even perfectly aligned cells would sum to activity with a six-fold component of only 0.003% of its mean (Fig 1C, Supp Fig 5). Doeller et al. (2010) reported a significant circular correlation between preferred direction (modulo 60°) and grid orientation in 18 conjunctive grid cells from eight rats (r = 0.29, p = 0.003), but our data did not show a significant correlation (n = 890 conjunctive cells meeting their criterion, r = 0.001, permutation test, p = 0.16; Fig 2D). The difference could be due to the number of cells and animals or to the recording site and method.

**Figure 2.**
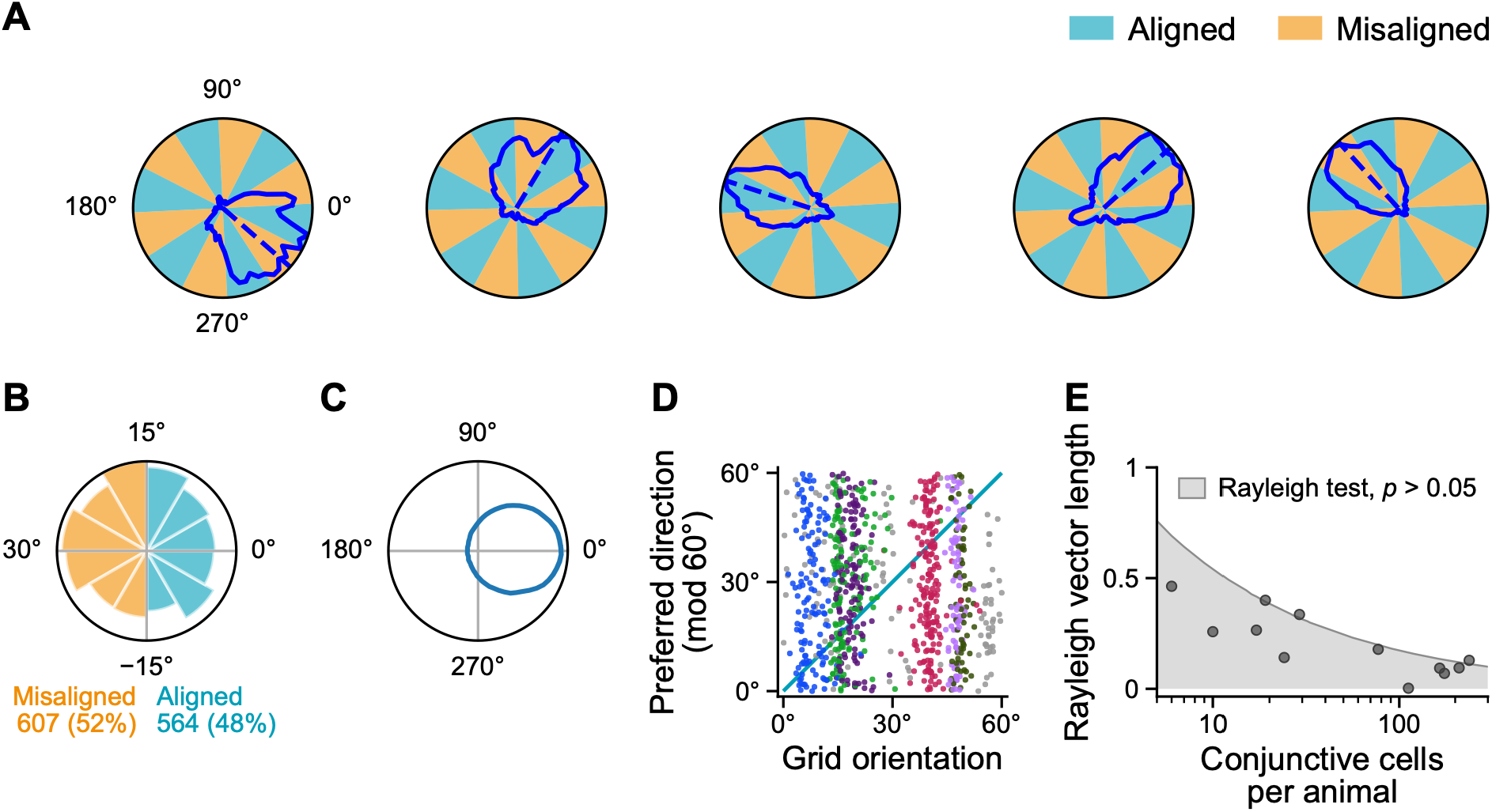
Conjunctive cells are broadly tuned and not aligned with the grid axes. **A.** Directional tuning curves (blue line) of five example conjunctive cells; dashed line, preferred direction; shading, aligned (blue, within 15° of a grid axis) and misaligned (orange) sectors. **B.** Polar histogram of the deviation of each cell’s preferred direction from the nearest grid axis (0°): 564 cells (48%) aligned, 607 (52%) misaligned. **C.** Mean tuning curve: each cell’s curve, divided by its mean and centred on its preferred direction (0°), averaged across animals. Median circular standard deviation across all 1,171 cells, 73°. **D.** Preferred direction modulo 60° against grid orientation for the 890 grid cells meeting the directional criterion of Doeller et al. (2010), plotted as in their Fig 1b. Cells with a preferred direction on a grid axis would fall on the diagonal line (teal); both axes are circular. Colours, the six animals with the most cells (7CC cells); grey, other animals. Each animal forms a vertical band because its cells have similar grid orientations. **E.** Clustering at 60° intervals within animals. Rayleigh vector length of six times each cell’s preferred direction (0, no clustering at 60° intervals; 1, perfect clustering at 60° intervals) against number of cells (12 animals of Vollan et al. (2025), 1,082 cells). Grey band, Rayleigh test p > 0.05; no animal is significant after Holm correction.

Conjunctive cells could still produce HexMod if their preferred directions clustered at 60° intervals around an angle other than the grid axes. As grid cells within each animal could cluster at different angles, we tested separately each of the 12 animals of Vollan et al. (2025) with at least five conjunctive cells (Fig 2E). Three animals showed clustering at 60° intervals with p < 0.05 (Rayleigh test), but none remained significant after multiple comparison correction (Holm-corrected p ≥ 0.23, n=12). Given the broad tuning, such slight clustering would add almost no six-fold modulation (Fig 1C), so conjunctive cells are unlikely to contribute to HexMod.

### Firing is suppressed shortly after visiting a grid field

Next, we set out to test the adaptation hypothesis using data from 3,938 grid cells from 32 animals (112 sessions), including both those with and without conjunctive directional tuning. Firing rate adaptation predicts that a cell should be less excitable shortly after recent activity, with excitability recovering over time. To test this prediction, we delineated each cell’s grid fields and measured the time elapsed since the most recent visit to any grid field (*t_prev_*) (Fig 3A,B). We then estimated the mean firing rate for different values of *t_prev_*, while controlling for location- and speed-dependent firing using a Poisson multiplicative model (see Methods). In line with an adaptation account, firing rate increased significantly from short to long values of *t_prev_* (Fig 3C; repeated one-way ANOVA F(3,11811)=375.8, p<0.001, followed by post hoc t-tests: all pairs p < 0.001 except 2–3s versus > 3s, p = 0.12).

**Figure 3.**
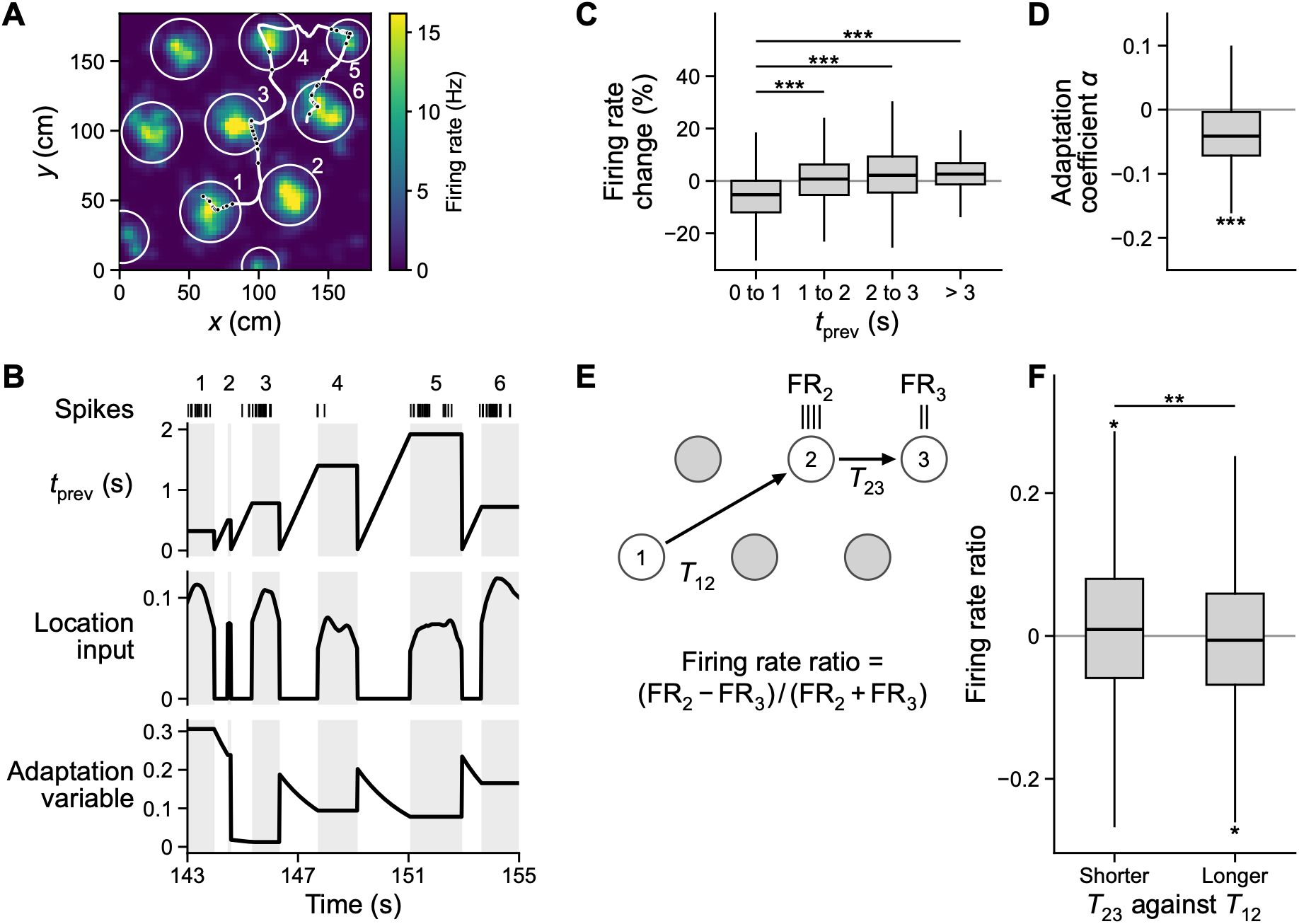
Evidence of firing rate adaptation in grid cells. **A.** Rate map of one grid cell is shown with an example12 s trajectory (white line), spike locations (black dots) and grid fields (white circles); fields are numbered in order of visit. **B.** Time courses along the trajectory in A. Grey shading, period inside grid fields, numbered as in A; ticks (Spikes), time bins with spikes. Top, time since the previous field visit (t_prev_). Middle, location input: a Gaussian of the distance from the field centre, weighted by the field’s peak firing rate. Bottom, adaptation variable: exponential decay of t_prev_, weighted by the integrated location input. In this example, adaptation is expected to be maximal after exiting field 1. **C.** Firing rate (FR) change (%) as a function of when the previous field was visited (t_prev_). In line with the adaptation hypothesis, FR was significantly lower for shorter t_prev_(n = 3,938 cells, ANOVA, and post-hoc t-test p<0.001). **D.** Adaptation coefficient derived from a parametric adaptation model was significantly negative across cells, meaning lower FR after recent activity. **E**. The adaptation hypothesis predicts a firing rate reduction from field 2 (FR_2_) to field 3 (FR_3_) during three consecutive field visits when time interval between field 2 and 3 (T_23_) is shorter than the interval between field 1 and 2 (T_12_). **F.** Firing rate ratio for sequences of three field visits in which the second interval was shorter (T_23_ < T_12_) or longer (T_23_ > T_12_) than the first. Boxes show the median and interquartile range and whiskers extend to the most extreme values within 1.5 interquartile range (same convention used in all figures).

Next, we tested a more realistic model of adaptation, in which a cell’s excitability depends not only on the time since recent activity but also on how strong that activity was. For instance, excitability should be lower after the animal spent a long time near the centre of a grid field than after a brief pass through its edge (e.g. field 1 versus field 2 in Fig 3A). We therefore defined an adaptation variable (Fig 3B, bottom) as the product of two terms: the expected neural activity due to the most recent field visit, estimated by a Gaussian function of the animal’s distance from the centre of the grid field, weighted by the field’s firing rate (Fig 3B, middle); and an exponential decay of the time since that visit, with a time constant of 2s (see Discussion for the choice of time constant). We then modelled each cell’s firing rate as the product of location-dependent, speed-dependent and adaptation terms, with Poisson noise (see Methods). The adaptation coefficient was significantly negative across cells, meaning lower firing after stronger recent activity (Fig 3D; t(3937) = −41.1, p < 0.001, Cohen’s d = −0.65).

### Firing rate adaptation across successive field visits

The preceding analyses showed the existence and time course of firing rate adaptation using entire recordings which include periods when an animal was located between grid fields. We next compared firing rates within consecutively visited fields as a function of the recent visit history - more closely matching the mechanism proposed to underlie HexMod, which concerns firing in one field shortly after visiting another field. For each cell, we identified sequences of three consecutive field visits (see Methods) and compared firing rates in the second (*FR*_2_) and third fields (*FR*_3_) (Fig 3E). Under adaptation, firing should be lower in whichever of these two fields was entered sooner after the preceding field visit. We summarised this using a normalised firing rate ratio, calculated as the difference between the rates in the second and third fields divided by their sum. The ratio should be positive (higher firing in the second field) when the interval between the second and third fields was shorter than the interval between the first and second fields (*T*_23_ < *T*_12_), and negative in the opposite case. Consistent with this prediction, the firing rate ratio was positive when the third field was reached after a shorter interval than the preceding transition (T_23_ < T_12_: mean 0.008, t(3575) = 2.53, p = 0.012, Cohen’s d = 0.04) and negative when it was reached after a longer interval (T_23_ > T_12_: mean −0.006, t(3575) = −2.10, p = 0.036, d = −0.04), giving a paired within-cell difference of 0.014 (t(3575) = 2.90, p = 0.004, d = 0.05; n = 3,576 cells with sequences in both conditions; Fig 3F). Adaptation is therefore visible at the level of individual transitions between consecutive field visits, the situation in which it could produce HexMod.

### Firing rate adaptation is a robust and stable feature of grid cells: control analyses

Next, we carried out several control analyses to assess the robustness of the firing rate adaptation effect. First, we tested its internal consistency by dividing each recording into even and odd minutes. The adaptation coefficient was significantly negative across cells in each half of the recording (Supp Fig 1A). Furthermore, adaptation coefficients were significantly correlated between the two halves, indicating that some cells consistently show stronger adaptation than others (Supp Fig 1B).

Second, we tested whether the lower firing observed shortly after leaving another grid field (short *t_prev_*) could instead be explained by prospective coding, in which firing is tuned slightly ahead of the animal’s current location (Chaudhuri-Vayalambrone et al., 2023a; Csordas et al., 2024; De Almeida et al., 2012; Ouchi & Fujisawa, 2024). With prospective firing, a cell starts firing before the animal enters a field and stops before it leaves. Positions near the edge of a field are therefore associated with higher firing on entry and lower firing on exit. Because short values of *t_prev_* necessarily occur shortly after leaving a field, prospective coding can produce lower firing at short *t_prev_*, mimicking adaptation. To examine this possibility, we shifted spike times relative to position (−400 to +400ms in 20ms steps) and selected the time shift that gave the highest spatial information (Supp Fig 2A,B). The median temporal shift was 40ms across cells (Supp Fig 2C). We then computed the adaptation effect after controlling for prospective coding, by shifting spike times accordingly, and found that it remained highly significant albeit slightly reduced (Supp Fig 2D,E). Hence, the adaptation effect exists in addition to any impact of prospective coding.

**Supplementary Figure 1.**
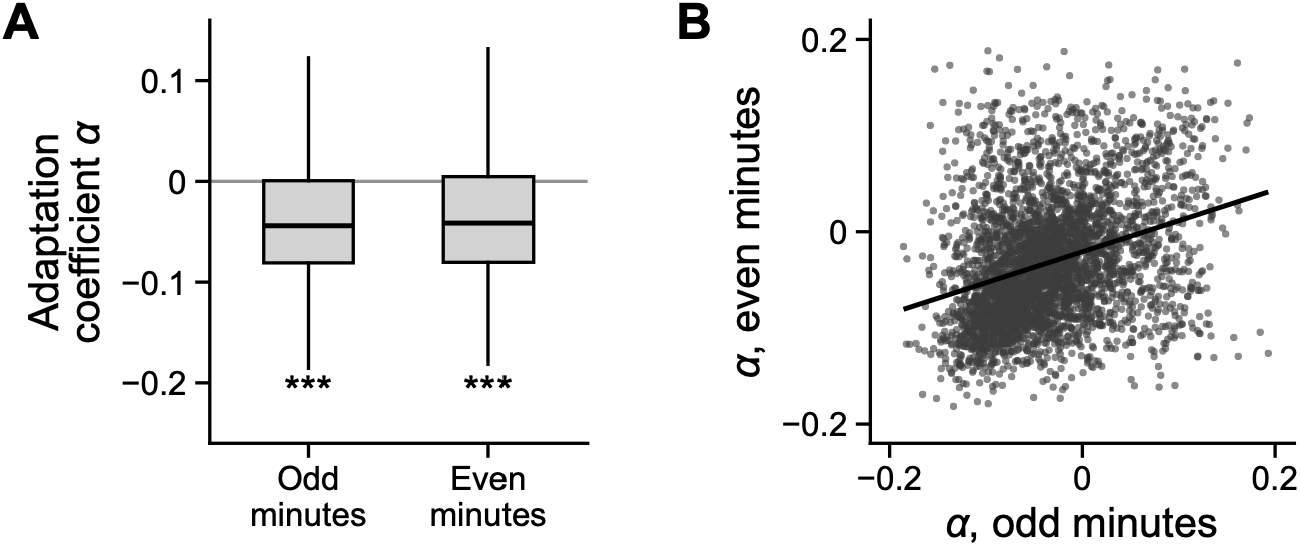
Adaptation is a robust and stable feature of grid cells. Each trajectory was divided into halves (odd and even minutes) and we applied the parametric adaptation model in each half of the data. **A.** The adaptation coefficient was significantly negative in each half. **B.** Adaptation coefficient α was significantly correlated between the two halves (r(3936)=0.31, p<0.001). It means that some cells consistently show strong adaptation (more negative α) and some cells do not (zero or positive α).

**Supplementary Figure 2.**
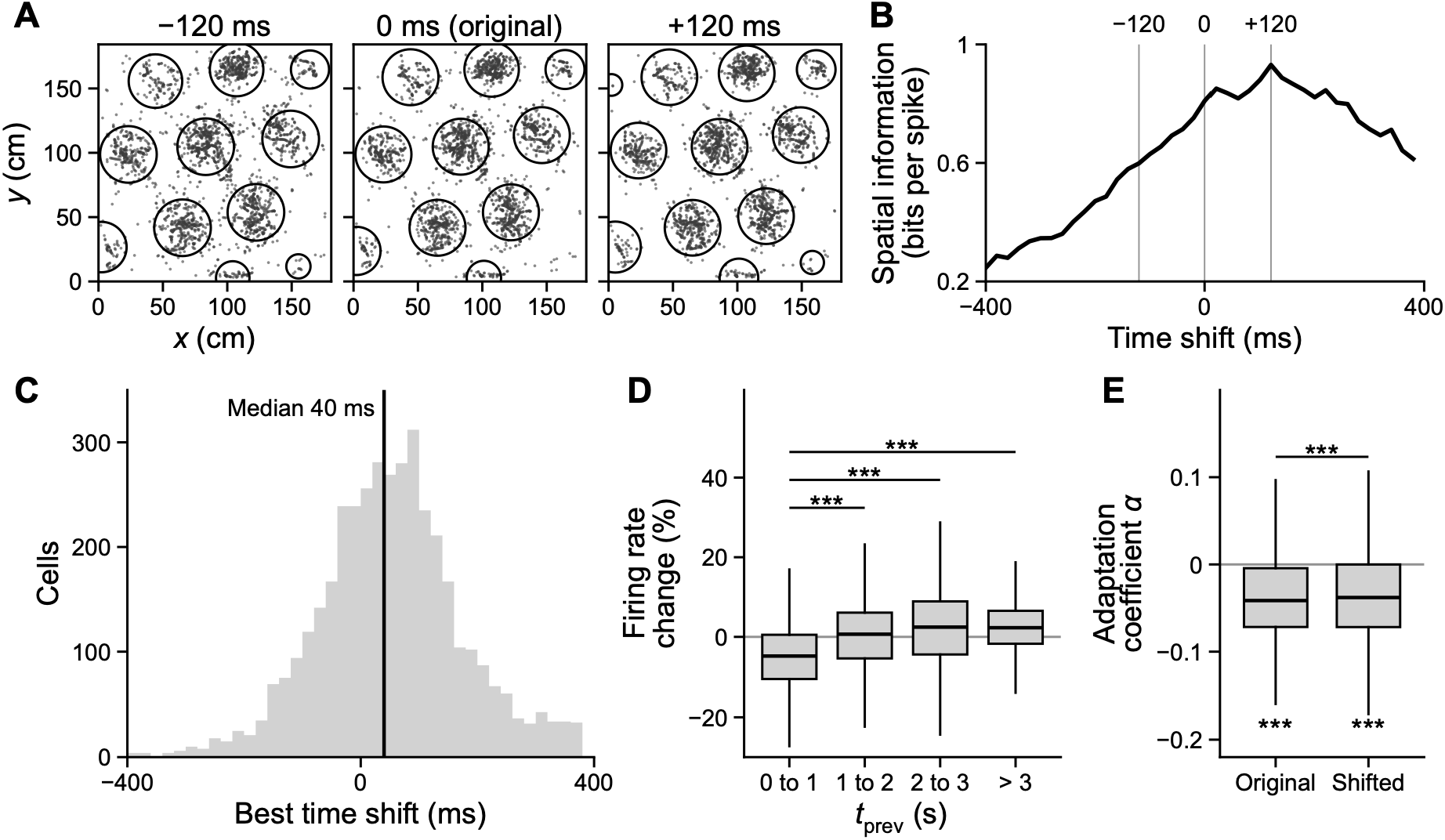
Adaptation remains significant after controlling for prospective coding. First, we shifted spike times relative to the behavioural trajectory from −400 to 400 ms and found the temporal shift that gave the highest spatial information for each cell. **A**. Spike positions (grey dots) of an example cell and its delineated grid fields (black circles), with spike times shifted by −120 ms, 0 ms (original) and +120 ms relative to position; axes in cm; the fields were delineated separately at each shift. **B**. Spatial information of the example cell in A (bits per spike) against the time shift; the grey lines mark the three shifts shown in A. Spatial information was highest at +120 ms, meaning that the cell encoded locations ahead of the animal (prospective coding). **C.** The distribution of optimal time shifts across cells. Black line shows the median shift (40 ms). **D**. After shifting spike times accordingly, firing rates (FR) remained lower after more recent visit to a grid field (i.e. shorter t_prev_), supporting the adaptation hypothesis (F(3,11259)=294.7, p<0.001). **E**. Adaptation coefficient derived from a parametric adaptation model also remained significantly negative after shifting spike times, albeit slightly lower after shifting (e.g. less negative, t(3753)=−4.13, p<0.001)

### Grid-aligned movement and time between field visits in tortuous trajectories

Previous sections showed that grid cells show a robust firing rate adaptation (i.e. reduced firing after short inter-field intervals), and we now ask whether grid-aligned movements cause more adaptation than misaligned movement - a prerequisite for HexMod. In straight trajectories as in Fig 1D, aligned movements are associated with shorter inter-field interval, but this relation is not guaranteed if movement trajectories are tortuous. To examine this in rodent data, we measured the temporal proximity between grid field visits (defined as the exponential decay of time since the last field visit with a time constant of 2s) as a function of whether movement direction in the current field was aligned or misaligned (Fig 4A,B). We found that mean temporal proximity was significantly higher (i.e. shorter *t_prev_*) during aligned than misaligned movement (t(3937)=11.1, p<0.001, Fig 4C), and that this difference was more prominent for small (<62cm) than medium (62-86cm) or large (>86cm) scale grid cells (F(2,3935)=7.1, p<0.001, Fig 4D). Although this difference was highly significant, the mean temporal proximity for aligned movements was only about 1% higher than for misaligned movements (Fig 4C).

**Figure 4.**
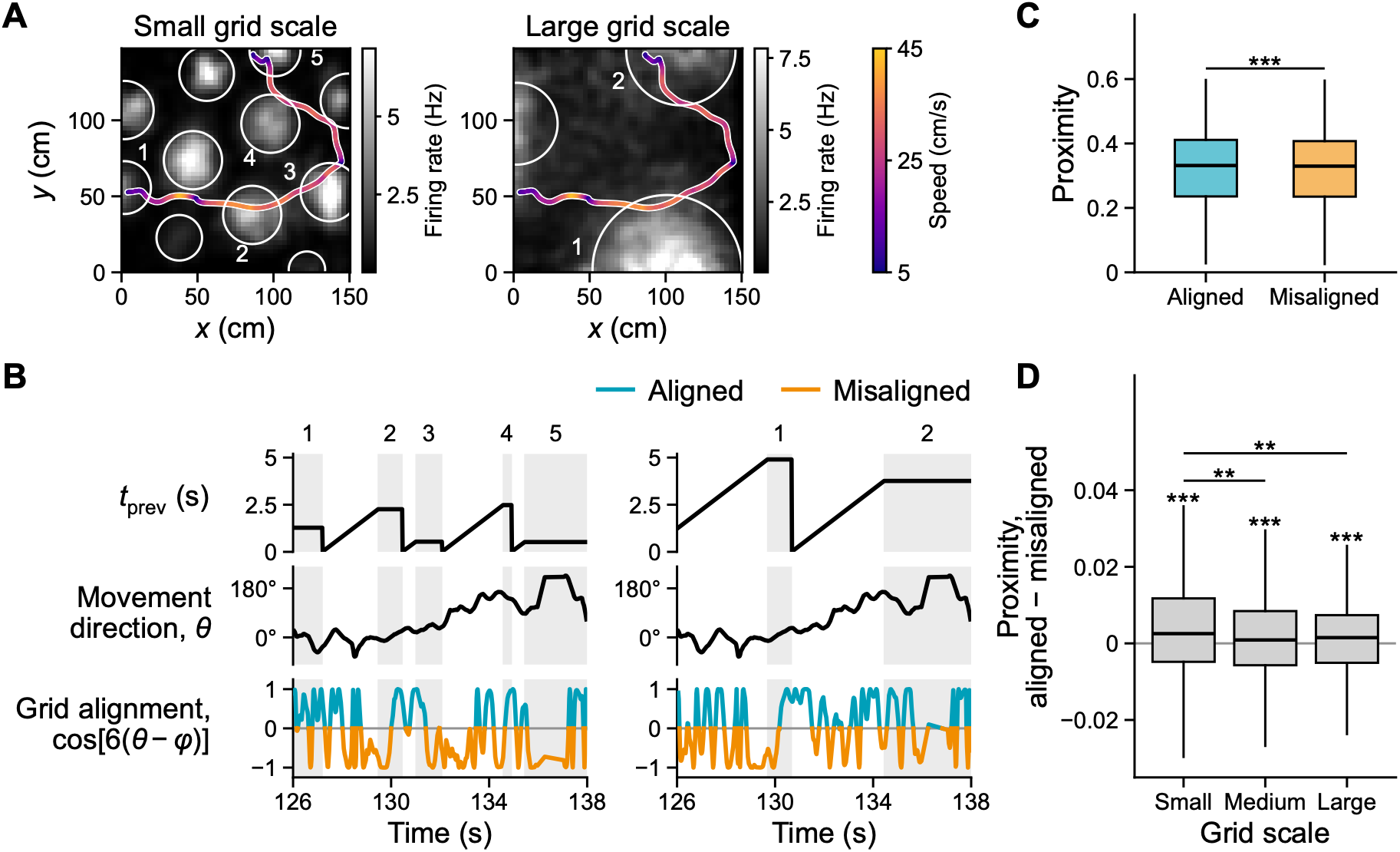
Grid alignment of movement and the time taken to move between grid fields in a tortuous animal trajectory. **A.** An example 12 s trajectory (coloured line) is overlaid on the rate map of both a small (left) and large (right) scale grid cell from the same recording. Colour indicates running speed. **B.** Time since the previous field visit (top), instant movement direction (middle), and grid alignment (bottom) for the trajectory in A are shown for the small (left) and large (right) scale grid firing patterns. Due to the frequent change of movement direction, grid-alignment is a weak predictor of time since the previous grid firing field visit (t_prev_). Grid alignment is measured as cos[6(θ − φ)], the cosine of six times the current movement direction relative to grid orientation, so that positive values indicate aligned (blue) and negative values misaligned (orange) movement directions. **C**. We defined proximity as the mean value of exponentially decaying time since the last field visit - exp(−t_prev_/τ) - for each time point during the aligned or misaligned movement inside the subsequently visited field for each cell. High proximity values therefore indicate a short interval between each grid firing field visit, and vice versa. Mean proximity was significantly higher during aligned versus misaligned movement periods (t(3937)=11.1, p<0.001), but the difference was small (only 0.7% higher for aligned movements). **D.** Proximity differences between aligned and misaligned movement periods are larger for small than medium or large-scale grid cells.

This weak association between movement direction and inter-field interval is due to tortuous trajectories, variable movement speed, and imperfections in the hexagonal geometry of grid firing patterns. While an animal is traversing a single grid field (e.g. field 2 in Fig 4A left panel), a slight change in movement direction (e.g. of 30°) within the field results in a complete transition from misaligned to aligned movement, rendering the grid alignment of current movement a poor predictor of time between fields. The association between grid alignment of current movement and time since the previous field visit is even weaker for large-scale grid cells, because there is a correspondingly longer time window within which the animal can change its direction both within and between grid fields (Fig 4A, right).

### Hexadirectional modulation depends on adaptation and trajectories

We next asked whether rat grid cells during free-foraging conditions show HexMod, and whether it depends on the two conditions identified above: strong firing rate adaptation, and movement paths with more recent field visits for aligned than misaligned movement. For each cell we estimated the strength of HexMod as the six-fold coefficient β: the amplitude of a six-fold (60°-periodic) modulation of firing rate by movement direction relative to the cell’s grid orientation (the factor 1 + β cos[6(θ − φ)], with θ the movement direction and φ the grid orientation), estimated with a multiplicative Poisson model that controls for location and speed (see Methods). β is positive when firing is higher during aligned movement and negative when it is higher during misaligned movement (30° offset); the adaptation hypothesis predicts negative values (Fig 1D). HexMod was not apparent when all cells were included (t(3937)=−0.2, p=0.8, Fig 5A), as expected from the weak association between the movement direction inside a field and the time since the previous field visit on tortuous trajectories (Fig 4C). We then restricted the analysis to the third of cells whose fields were visited in quicker succession during aligned than misaligned movement, that is, the top third of the aligned-minus-misaligned proximity difference (see Methods). Among these cells, those with the strongest adaptation (n=435) showed a weak but significantly negative HexMod β (t(434)=−2.8, p=0.005, Cohen’s d=−0.14), whereas mean HexMod β was near zero in cells with weak (t(429)=0.08, p=0.9) or no adaptation (t(447) = 0.8, p=0.4) (Fig 5B). In contrast, cells with strong adaptation but in the bottom third of the proximity difference did not show significant HexMod (t(442)=1.4, p=0.2, Fig 5C), reaffirming that both adaptation and trajectory conditions are necessary for HexMod.

**Figure 5.**
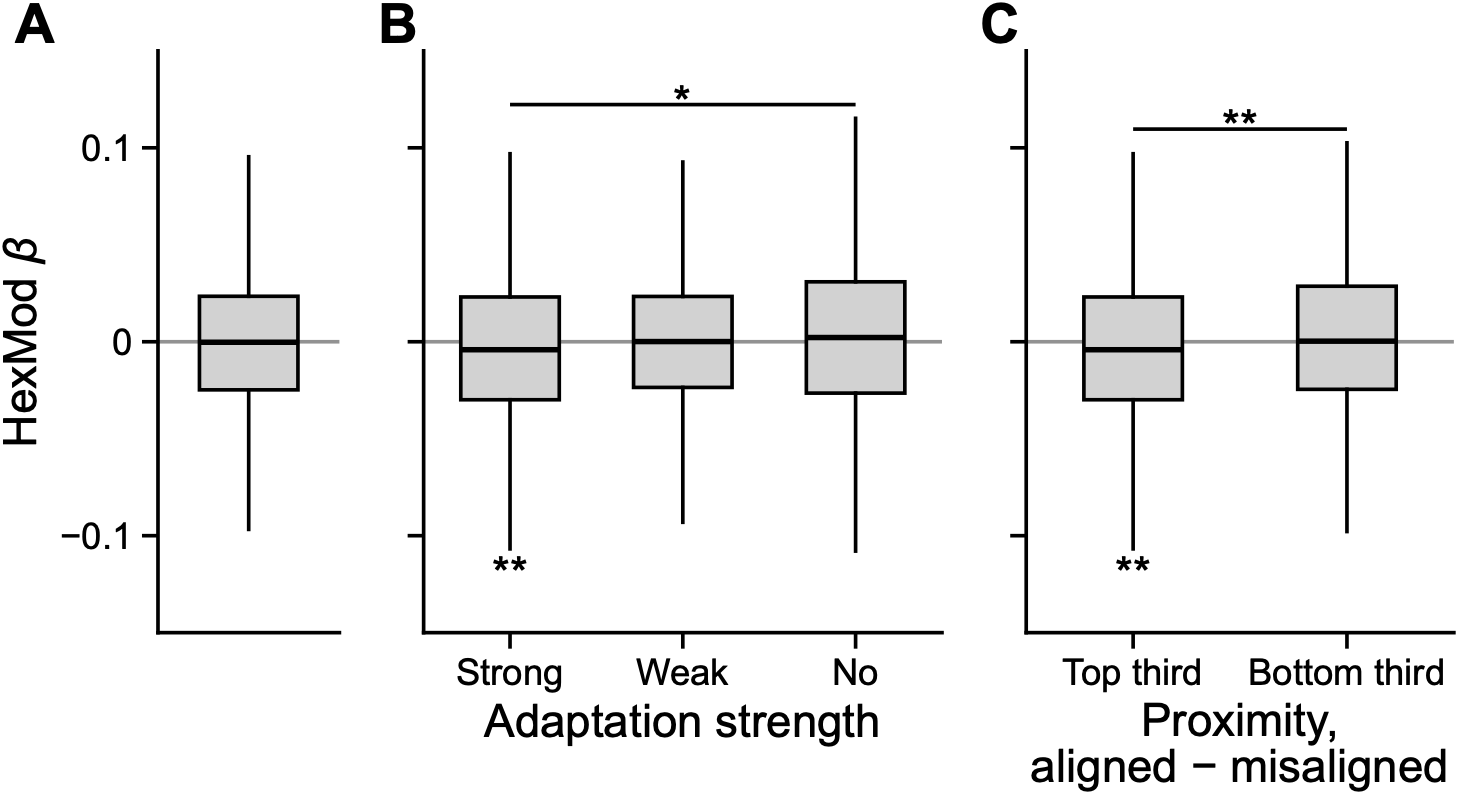
Both adaptation and trajectory conditions are necessary to produce hexadirectional modulation. **A.** When all grid cells are included, HexMod β was not significantly different from zero (t(3937)=−0.2, p=0.8). **B**. When restricting the analysis to cells with higher proximity for aligned movement (top 1/3 of cells), cells with strong adaptation showed a significant negative HexMod (t(434)=−2.8, p=0.005, Cohen’s d=−0.14), but cells with weak (n=430) to no (n=448) did not. **C.** Among cells with strong adaptation, only cells with higher proximity for aligned movement (top 1/3) showed a significant negative HexMod (identical to the strong group in panel B) whereas cells in the bottom third of the proximity difference did not differ from zero (t(442)=1.4). The two proximity groups differed significantly (t(876)=−3.0, p=0.003).

### Simulating grid cell population activity for human experiments: hexadirectional modulation and grid scale

The rat data suggest that adaptation can produce HexMod, but that tortuous foraging paths largely mask it. Human neuroimaging experiments typically use straighter paths, for example navigation to goal locations in virtual reality. We therefore simulated grid cell population activity during human-like straight trajectories, to ask whether HexMod would be detectable under these conditions and how it depends on grid scale. For each of 30 virtual participants we simulated two separate grid modules, one small-scale (grid scale 40% of the environment width) and one large-scale (70%), and analysed each module separately (Fig 6B,C). Each module contains 10,000 cells approximating the number of grid cells in a single 2.5 mm voxel of functional magnetic resonance imaging (fMRI) (see Methods). The participants moved in straight lines from random start locations to hidden goal locations (Fig 6A). Crucially, these less tortuous movement trajectories generate a large difference in inter-field intervals between aligned and misaligned movements, especially for the small-scale grid module (Fig 6D, cf. Fig 4C with tortuous animal trajectories). We modelled the firing rate of each grid cell as a product of location-dependent firing and firing rate adaptation that depends on the time since the previous field visit (see Methods). The summed firing rate of all cells can be viewed as a noise-free proxy for a macroscopic neural measurement, e.g. blood-oxygen-level-dependent (BOLD) fMRI signal in the entorhinal cortex. Our simulated grid cell firing rate maps had spatial imperfection and inhomogeneous firing rates across fields like real grid cells, because deviations from the perfect hexagonal lattice can also affect the HexMod (Supp Fig 3).

**Figure 6.**
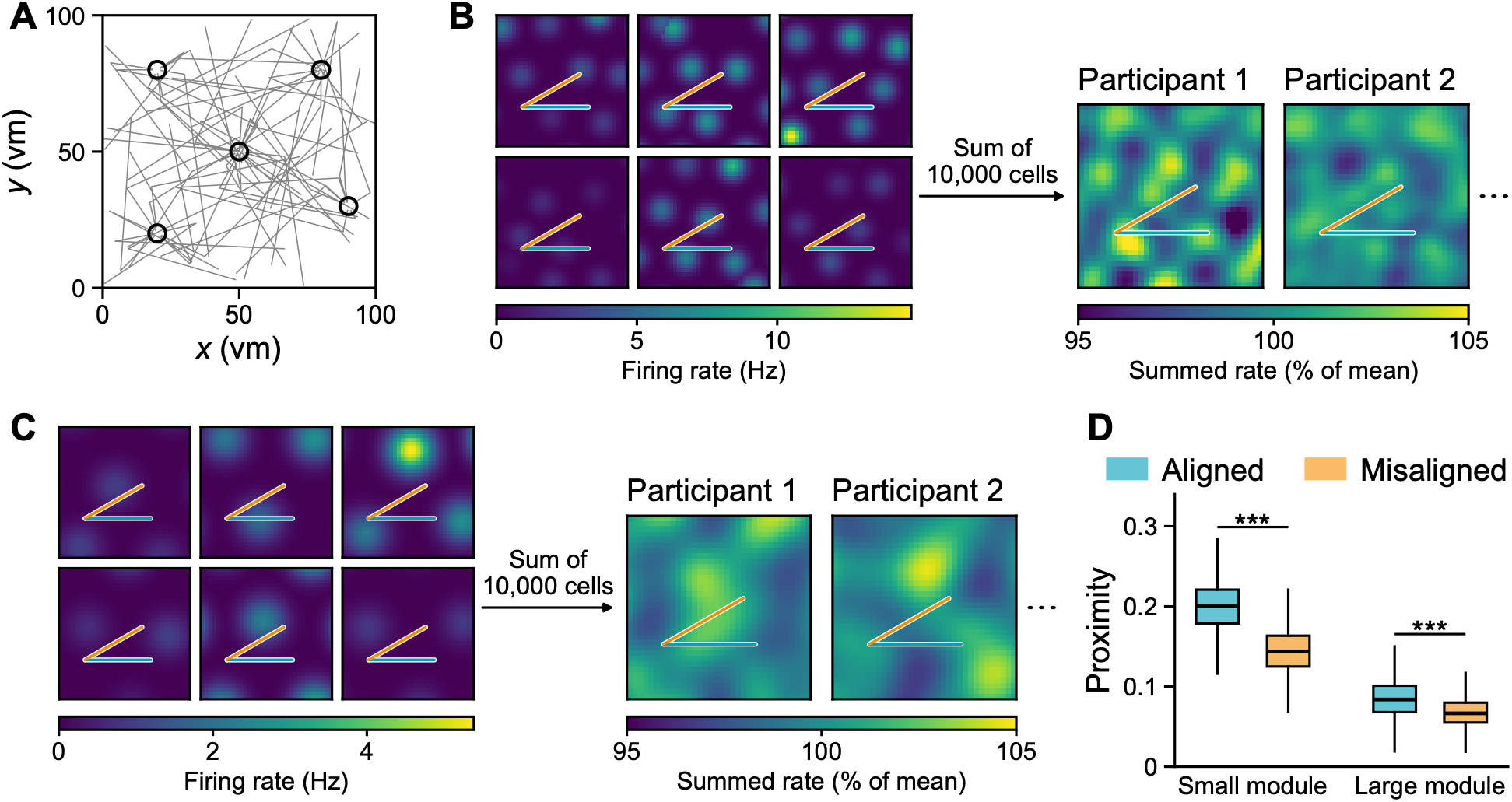
Simulation of grid cell population activity in a typical human neuroimaging experiment. **A**. Example trajectories for an object-location memory test. The simulated participants moved directly from random start locations to hidden object locations (black circles). As in typical desktop-based virtual reality navigation tasks in humans, movement speed is approximately binary (zero or maximal speed). **B**. Example small-scale grid module comprised of 10,000 grid cells with consistent orientation and scale (40% of environmental width) and random offsets. We applied a small random jitter to the location of firing fields and varied peak in-field firing rates to match realistic rate maps from rodent data. The summed activity of each grid module therefore had a small residual location-dependent firing driven by the random offset of grid firing fields and differs for each of the 30 virtual participants. Blue and orange lines, example aligned (along a grid axis) and misaligned (30° from it) movement segments. **C.** Same as **B** for large-scale grid module (scale=70% of environmental width). **D.** Proximity (exponential decay of time since the last firing field visit with a time constant of 2s) was significantly higher for aligned than misaligned direction, and the effect size was larger for small-scale modules. Data from one example module is shown here.

**Supplementary Figure 3.**
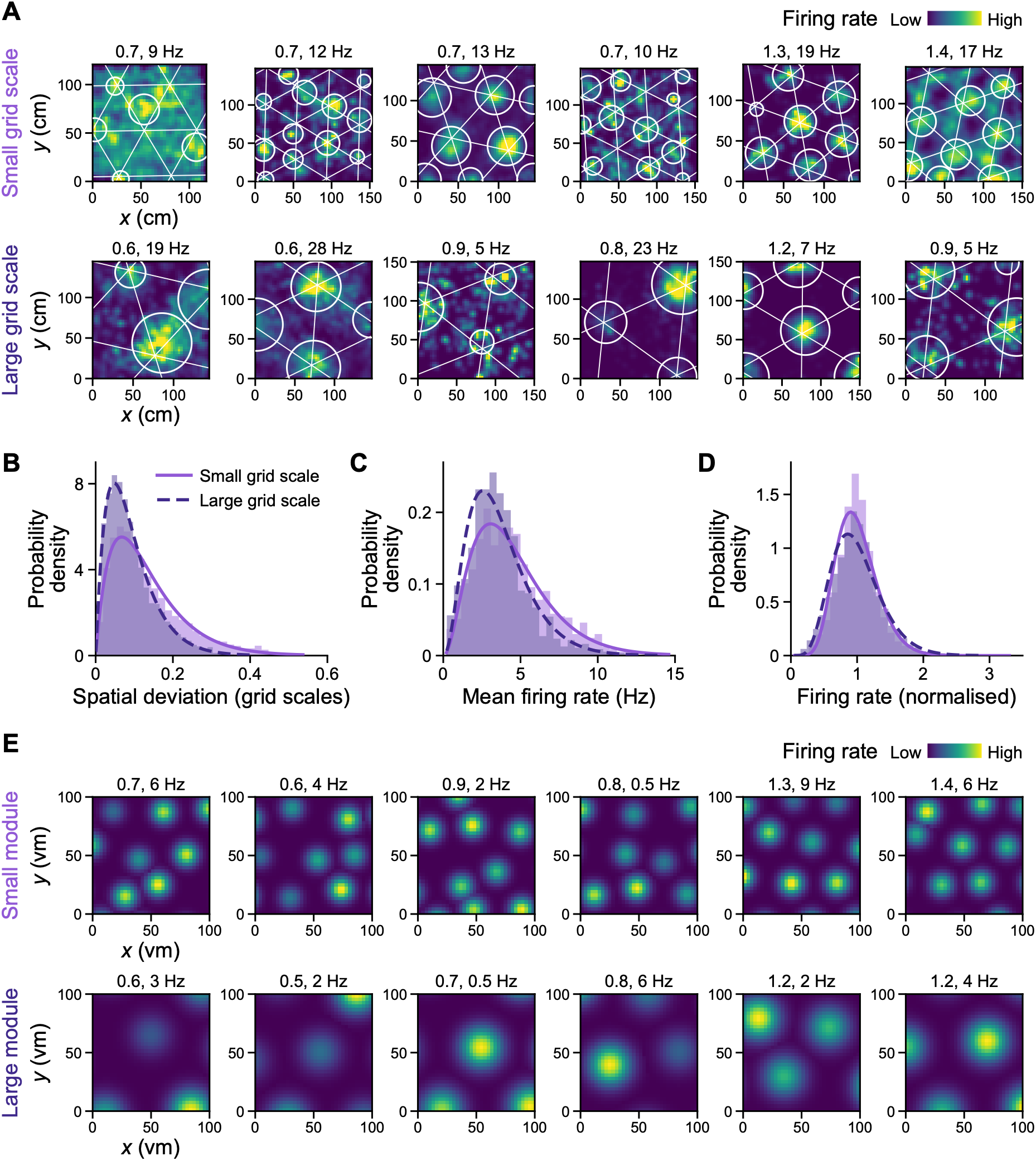
Grid cells deviate slightly from a perfect hexagonal lattice. **A**. Example rate maps of real grid cells from small-scale (< 0.4 * environment size) and large-scale (> 0.6 * environment size) modules. Bright colours indicate higher firing rates, white circles show delineated grid fields. White lines show the best fit triangular lattice to grid field centres. Gridness scores and peak firing rates are shown above each rate map. **B**. Distribution of spatial deviations, defined as the distance between the centre of grid fields and nearest triangular vertex, normalised by grid scale, fit by a gamma function. Small scale: shape=2.0, scale=0.07; Large scale: shape=2.1, scale=0.04. **C**. Distribution of mean firing rates across grid cells. Small scale: shape=3.2, scale=1.4; large scale: shape=3.3, scale=1.1. **D**. Distribution of normalised firing rates within cells. Small scale: shape=10.3, scale=0.1; large scale: shape=7.1, scale=0.1. **E**. Example firing rate maps of simulated grid cells with spatial deviation and firing rate variability across fields.

First, we checked the existence of HexMod by looking at mean firing rates during aligned and misaligned movement relative to the ground truth grid orientation. The small-scale grid module showed the expected pattern of lower firing rates for aligned versus misaligned movement trajectories due to firing rate adaptation (t(29)=−23.6, p<0.001, Cohen’s d=−4.4, Fig 7A). Mean firing rates were also lower for aligned versus misaligned movement trajectories in the large grid module, but the effect size was smaller (t(29)=−7.1, p<0.001, Cohen’s d=−1.3, Fig 7A).

**Figure 7.**
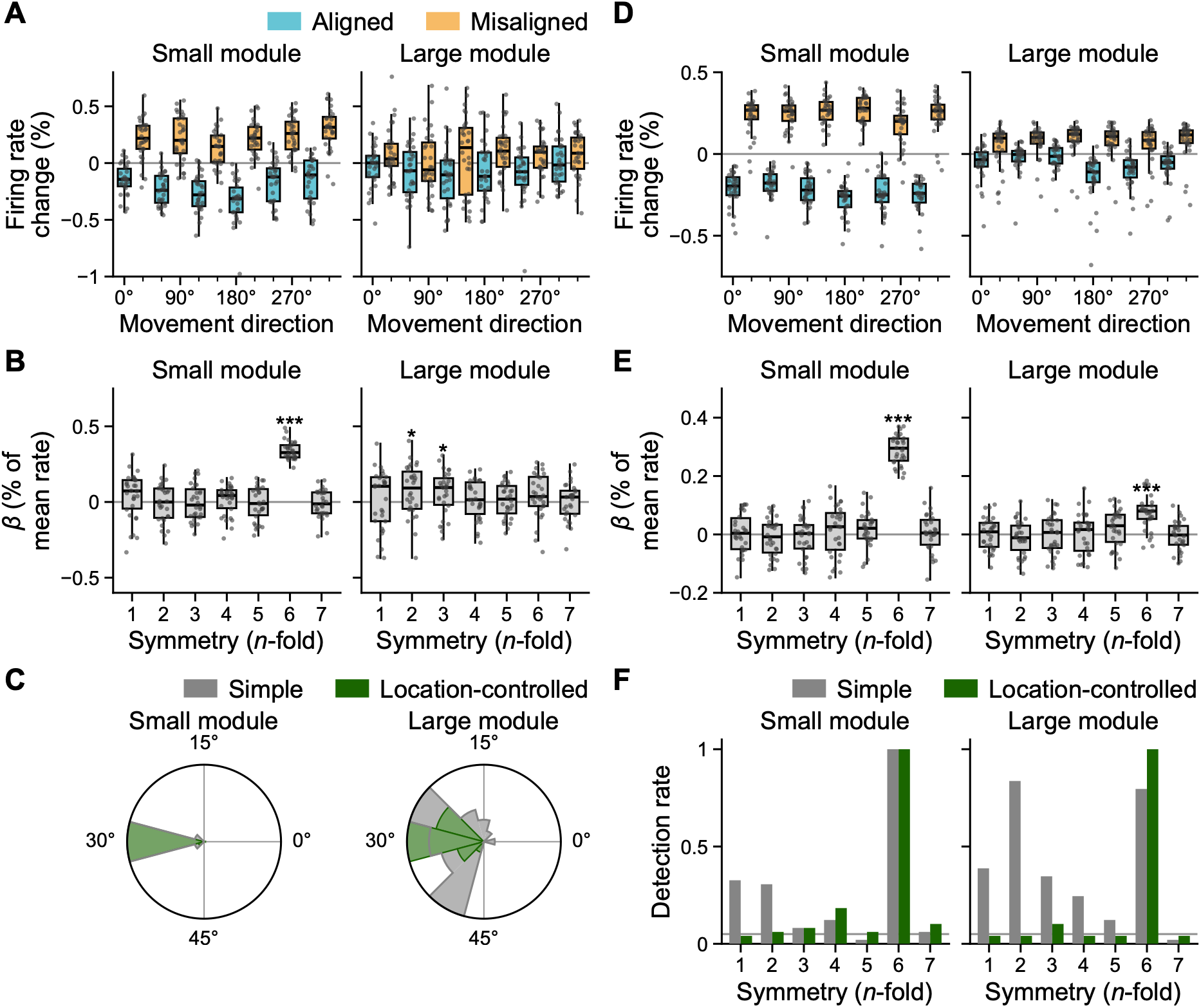
Detection of hexadirectional modulation is improved by a location-control estimation method. **A.** Normalised summed firing rate in small- and large-scale grid modules for each aligned and misaligned movement direction relative to the true grid orientation. Firing rate is lower for aligned (blue) than misaligned (orange) movement directions due to adaptation, especially for small grid modules. Individual data points (for 30 simulated participants) are overlaid. **B**. Cross-validated β coefficients for each n-fold symmetry, for small- and large-scale grid modules. In small modules, C-fold symmetry was significant, whereas 2- and 3-fold symmetry reached significance in large modules. **C**. Estimated orientation of peak activity from cross-validated HexMod analysis when the simple (grey) or location-controlling multiplicative (green) method was used. In small modules, the orientation is perfectly estimated as 30° relative to true grid orientation in both methods, showing overlapping grey and green. In large modules, estimated orientations from the simple method are more variable, while those from the multiplicative method are tightly clustered to the correct orientation. **D**/**E**. same as **A**/**B** with location-controlled estimation method. **F**. Detection rate for n-fold symmetry when the simulated experiment with n=30 participants was repeated multiple times. Compared to the simple method (grey), the location-controlled method (green) suppressed the spurious finding of low-fold symmetry and improved C-fold symmetry detection in both small- and large-scale grid modules, with the benefit being more evident in large scale.

Next, we applied the standard HexMod analysis method for neuroimaging data, which searches for consistent N-fold directional symmetry (i.e. cos[*N*(*θ* − *ϕ*)], where *θ* is movement direction and *φ* is the grid orientation of higher firing) (Doeller et al., 2010). This method first estimates the grid orientation from part of the data and then tests whether the remaining data follows the predicted N-fold cosine modulation relative to the estimated grid orientation. The significance of HexMod is then tested at the group level on the resulting regression coefficients. For small grid modules, six-fold symmetry was prominent (Fig 7B) and the estimated orientation of peak activity (*φ*) was tightly clustered around 30° relative to the true grid orientation (Fig 7C; the 30° offset showing that HexMod is negative rather than positive, consistent with firing rate adaptation). In contrast, large grid modules showed significant 2- and 3-fold symmetry, rather than the expected six-fold (Fig 7B), and the estimated orientation of peak activity was more variable (Fig 7C, grey).

There are multiple potential explanations for the weaker HexMod observed in large grid modules. First, two firing fields are rarely visited in quick succession, even during aligned movement, due to the large spacing between fields (see Fig 6C). Second, inhomogeneous firing rates across grid fields and spatial deviations from a perfect hexagonal lattice result in spatial modulation of summed firing rates, even across a large number of grid cells, such that population activity shows up to ±5% variation across the environment (Fig 6C). Summed rate maps are spatially smooth and smoothness scales with grid scale, forming larger elevated or depressed regions of activity. When a trajectory passes these regions in opposite directions, a strong 2-fold symmetry can occur and mask a weaker six-fold symmetry.

### Improving hexadirectional modulation detection by controlling for spatial variation

To mitigate the spatial modulation of population activity due to inhomogeneous grids, which can confound HexMod, we modelled the summed activity as the product of a spatial map and a directional profile (a location × direction multiplicative model; see Methods). The directional profile then measures how activity varies with movement direction once the uneven sampling of locations in each direction has been accounted for. This method can be applied to human neuroimaging data. With this method, firing rate differences between aligned and misaligned runs are amplified (Fig 7D), and only 6-fold symmetry becomes significant in the cross-validated method for both small- and large-scale modules (Fig 7E). Estimated orientation of peak activity also became more tightly clustered relative to the true grid orientation in the large grid module (Fig 7C, green). We repeated the simulation of 30 virtual participants 50 times using different random grid offsets, peak firing rate, and trajectories, to test how often 6-fold and other symmetries become significant when using the standard simple method and location-controlled model. When the simple method was applied, 6-fold symmetry was most prominent in small modules, whereas 2-fold symmetry was found more often than 6-fold symmetry in large modules (Fig 7F, grey). Using the location control method, only 6-fold was significant (Fig 7F, green). Recently, a method to reduce the false positive inference of 6-fold symmetry due to 1-,2-,3-fold symmetry was proposed (Almog et al., 2026). In contrast, our new control method reduces a false negative inference and could potentially improve the sensitivity of detecting grid cell activity.

### Hexadirectional modulation strength depends on movement speed

It has previously been demonstrated that HexMod is stronger during faster movement periods (Doeller et al., 2010). The adaptation hypothesis offers an explanation for this. For aligned movement, the time between consecutive grid fields is about one grid scale divided by running speed. For misaligned movement it is about √3 (≈1.7) times longer, because the nearest field in a misaligned direction is further away (Fig 8A). Because adaptation decays exponentially with this time, firing is suppressed more during aligned than misaligned movement (Fig 8B). This difference is small at low speeds, when both intervals are too long compared to the adaptation time constant, and at high speeds, when both are short. It therefore peaks at an intermediate speed (Fig 8C). The optimal speed is proportional to the grid scale divided by the adaptation time constant, about 1.33 × scale/τ (see Methods), so small-scale modules are best detected at slower movement speeds and large-scale modules at faster ones.

**Figure 8.**
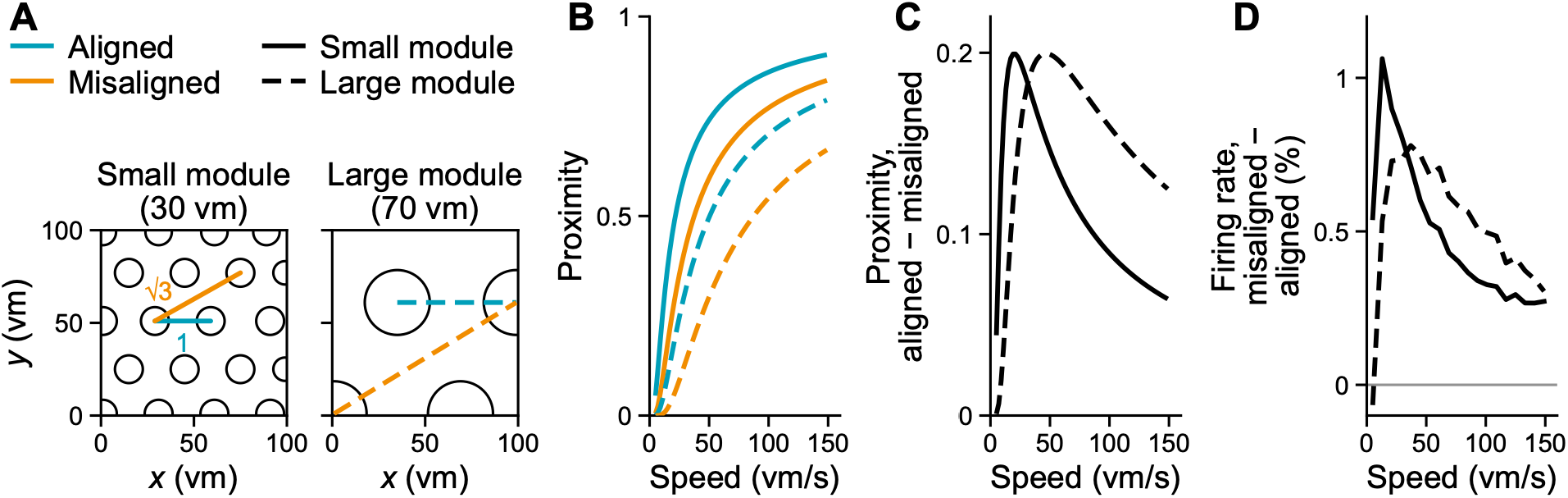
Prediction for the effect of speed on hexadirectional modulation strength. **A.** The time between the neighbouring field (aligned, blue) is one grid scale divided by speed, and the time between the non-neighbouring field (misaligned, orange) is 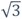 of the aligned. Grid fields are shown for a small and large grid cell. **B.** A proximity, exp(−time/τ), for aligned (blue) and misaligned (orange) monotonically increases with speed. Proximity is higher for smaller grid scales. Time constant τ is 2 s in this example. **C.** Proximity difference between aligned and misaligned movement (aligned − misaligned). It is maximal at a speed of about 1.33 × grid scale/τ; the firing rate difference produced by adaptation is proportional to it. **D.** Simulation of ideal grid cells in continuous long trajectories showed a speed-dependency comparable to that in C. Solid line: small scale, dashed line: large scale.

Simulation of ideal grid cell populations in simplified condition (a long continuous trajectory from a wall to a wall, see Methods) confirm the non-monotonic speed dependence on HexMod strength (Fig 8D). This finding is consistent with a previous simulation study which reported that the optimal adaptation time constant is roughly the grid scale divided by the speed (Bin Khalid et al., 2024). Of note, the effect of speed would be more complicated in a real experiment because the time between fields is not solely determined by the movement speed but also by non-movement periods (e.g. turning and inter-trial-intervals), changing of movement direction at boundaries, and imperfect geometry of grid fields.

## Discussion

We sought to adjudicate between two hypotheses for the neuronal origin of the hexadirectional modulation (HexMod) observed in human neuroimaging studies: directional tuning of conjunctive grid × head-direction cells, and firing rate adaptation. Across a large population of rat grid cells, conjunctive grid cells showed broad directional tuning that was not aligned with the grid axes, therefore unlikely to cause HexMod. Instead, grid cell firing was reduced after recent activity, consistent with firing rate adaptation and capable of producing HexMod. Critically, adaptation can produce HexMod only when trajectories are sufficiently straight for grid-aligned movement to correspond to more recent field visits. Free-foraging rat trajectories were highly tortuous, yielding little difference in adaptation between grid-aligned and misaligned movement and consequently weak HexMod. Our simulations show that a robust population-level HexMod emerges with the straighter trajectories typical of human neuroimaging paradigms. Furthermore, we proposed a method to improve the detection of HexMod by controlling for the residual spatial modulation of grid cell activity.

### Long straight trajectories are required to detect HexMod

HexMod can arise when sampling grid fields in rapid succession along aligned directions compared to misaligned directions causes greater firing rate adaptation. This condition occurs when the trajectory is straight over a minimal length of the inter-field distance (~ grid scale) with even sampling of movement speed across directions (Bin Khalid et al., 2024; Mei et al., 2026). Trajectories that do not meet this condition might be the reason for non-significant HexMod effect reported in some human fMRI studies (Kransberg et al., 2026). When a fixed length of trajectory is used, small scale grid cells can show stronger HexMod than large scale grid cells due to geometry, as observed in the rodent data and simulations here. Given the anatomical gradient of grid scale, we predict a stronger HexMod strength in dorsal (rodent) and posterior (human) entorhinal cortex (Brun et al., 2008). Of note, a lower signal-to-noise ratio in the anterior entorhinal cortex due to fMRI susceptibility artefacts is a potential confounding factor for future investigations.

### The temporal profile of adaptation and its relation to speed, grid scale and HexMod

Reduced neural activity in response to repeated stimulation can occur on multiple timescales (Benda, 2021). For HexMod, adaptation should occur at the timescale of inter-fields interval, which is grid scale divided by movement speed. We are unaware of previous studies that directly measures the adaptation of grid cells, and only a limited number of in-vitro studies examined stellate and non-stellate cells in the rat entorhinal cortex. These found a strong reduction in spikes within 200ms of large current injections, followed by gradual reduction at longer time scales, 2±1s (Alonso & Klink, 1993; Yoshida et al., 2013).

Our analysis provides the first evidence of adaptation on sub-second and multi-second timescales in grid cells in foraging rats (Figure 3C, Supp Fig 4); the interval between successive field visits in small grid modules (median 1.4s in our trajectories) falls within this range. We also looked at the adaptation timescale in terms of firing rate reductions as a function of the temporal proximity of the most recent grid field visit in more detail. We found evidence for two timescales, of approximately 0.3 and 2.6s, roughly consistent with the in-vitro findings (Supp Fig 4). For estimating our simpler one-timescale parametric model, we chose a timescale of 2s.

The adaptation effect interacts with running speed and grid scale to produce the HexMod (Figure 8), implying that there is an optimal running speed for a given experiment. For virtual navigation where experimenters can control the movement speed, we recommend a movement speed proportional to the target grid scale. For instance, fast (5.3m/s) or slow (2.7m/s) speed is ideal for detecting large (8m) or small (4m) modules, respectively.

**Supplementary Figure 4.**
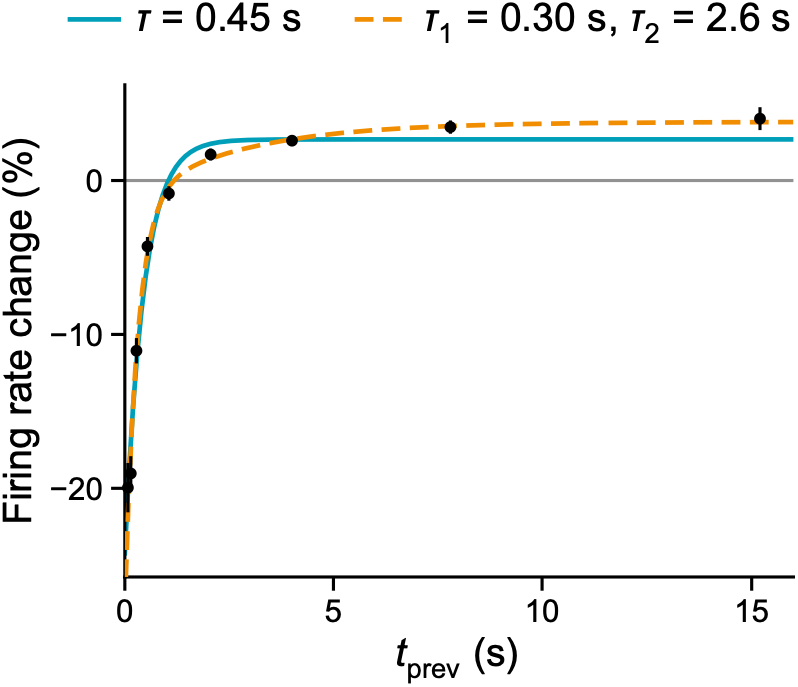
Timescales of firing rate adaptation. Firing rate as a function of time since the previous grid-field visit (t_prev_) was fitted with two models, a single exponential and a sum of two exponentials. Firing rate is expressed as percent change from each cell’s mean rate and averaged over cells (3,938 cells from 32 animals; 3,936 in the last bin, which two cells never reach). Points: mean and 95% CI in nine logarithmically spaced bins from 0.05 to 20 s. Teal: single exponential, τ = 0.45 s (95% bootstrap interval over animals 0.31 to 0.81 s). Orange, dashed: sum of two exponentials, τ = 0.30 s (0.21 to 0.56 s) and 2.6 s (1.3 to more than 10 s). Both models were fitted by least squares with each bin weighted by the inverse square of the standard error of its mean. The two-exponential model fitted better: χ^2^ = 21 with 4 residual degrees of freedom (9 bins minus 5 parameters) versus 84 with & (9 bins minus 3 parameters); ΔAIC = ΔBIC = −59; nested F(2, 4) = 6.1, p = 0.06. In 1,000 bootstrap resamples of animals, refitting both models each time, AIC preferred the two-exponential model in SS%.

### Ambiguity of grid orientation estimated from the traditional analysis

Human neuroimaging studies have estimated the grid orientation by assuming higher activity for aligned movements – opposite to the prediction from the adaptation hypothesis. Thus, according to our findings, the estimated grid orientation in these studies would correspond to 30° from the true grid axis. A conceptual 2D navigation task showed a stronger map-like distance code in the medial prefrontal cortex in aligned than misaligned trials (Qu et al., 2026), but this could be interpreted as a stronger code in misaligned trials. A visual search experiment found that grids are anchored to the border of square display (Julian et al., 2018). This finding remains valid despite the inverted grid orientation because the two borders are orthogonal in the square, so that an orientation at 7.5° from the North-South border would become an orientation at 7.5° from the East-West border. Researchers should be cautious about interpreting the orientation of the peak neuroimaging signal as aligned or misaligned to the actual grid firing patterns. An added complexity is the non-trivial relation between the neural and BOLD fMRI signal, with the possibility that higher BOLD signal might indicate lower neural firing (Ekstrom, 2010; Epp et al., 2026).

### Controlling for potential confounding factors when measuring HexMod

To cleanly measure the HexMod, location, speed, and direction should be evenly sampled but this is not always possible due to experimental constraints. Therefore, researchers can first check whether raw signal (e.g. BOLD) depends on potentially confounding factors such as location and speed using a simple regression model. If the answer is yes, they can apply the multiplicative model that separately estimates the effect of direction from other variables to uncover the hidden HexMod, as in our simulations. Controlling for uneven sampling of location and speed is even more relevant in cases where a significant N-fold symmetry is found for N < 6 when using a standard analysis pipeline (Sigismondi et al., 2024). Lower rotational symmetries could frequently occur from confounding factors of directional tuning (Almog et al., 2026), location, or uneven sampling of speed (e.g. faster speed along the border of a rectangular environment).

### Interpreting individual differences in HexMod

HexMod has shown dependence on task performance (Constantinescu et al., 2016; Doeller et al., 2010), age (Qu et al., 2026; Stangl et al., 2018), and risk of Alzheimer’s disease risk (Bierbrauer et al., 2020; Kunz et al., 2015). The individual differences in HexMod could be related to the integrity, maturity, and engagement of grid cells, but they can also reflect differences in the particular trajectories sampled, even when underlying grid cell population are similar. In addition, the electrical properties of neurons including afterhyperpolarisation – which in turn affects adaptation strength and therefore HexMod - can change during aging and neurodegenerative conditions (Wu et al., 2002).

### Limitations and future directions for non-invasive measurement of grid cells

First, we only included a population of pure grid cells from a single module in our simulation, but there are various spatially-tuned cell types in the entorhinal cortex which together outnumber the grid cells. Therefore, the HexMod strength measured from real fMRI data is expected to be significantly noisier than in our simulation, while detailed biophysical modelling of all cell types and neurovascular coupling is beyond the scope of the present paper. Second, we focused our rat data analysis on firing rate, rather than local field potential (LFP). HexMod of theta oscillations has been reported in human magnetoencephalography (MEG) (Convertino et al., 2023; Giari et al., 2023) and intracranial electroencephalography (EEG) (Chen et al., 2018; Maidenbaum et al., 2018). However, estimating theta power requires temporal windows of ~400ms, which may be too slow to capture the rapid directional changes of free-foraging rodents. Examining the origin of oscillatory HexMod signals in rodents will require carefully designed paradigms with constrained trajectories. Third, more comprehensive models for adaptation could be utilised. There are output-driven (e.g. calcium-activated potassium currents which inhibit spiking) and input-driven (e.g. short-term synaptic depression) mechanisms that could cause effects of adaptation (Benda, 2021; Gollisch & Herz, 2004). We mainly focused on the input-driven part, assuming that whenever a field is visited, inhibition accumulates regardless of actual spikes fired, for simplicity of analysis and modelling. We only modelled the adaptation effect from the latest field visit within short time scales, but there could be an accumulated effect of adaptation across multiple previous field visits.

### Optimising non-invasive measurement of neural population with firing rate adaptation

fMRI adaptation via repeated presentation of stimuli is a popular experiment technique for probing neural encoding from macroscopic signal, reviewed by (Barron et al., 2016). Indirect evidence of head direction cells and place cells in the human brain also came from fMRI repetition suppression, e.g. (Baumann & Mattingley, 2010; Doeller et al., 2010; Morgan et al., 2011). Ideally, the time-lag between the stimuli should match the temporal profile of adaptation in the targeted neural population. Most previous empirical data on neural adaptation comes from simple experimental paradigms using static images (Reber et al., 2019; Xiang & Brown, 1998). Our analysis is the first that quantitatively assessed the adaptation profile of grid cells in a naturalistic setting. It could be expanded to investigate the effects of adaptation of other cell types to improve the design and interpretation of other functional neuroimaging paradigms.

### Conclusion

Grid cells in foraging rats exhibit firing rate adaptation that can result in macroscopic hexadirectional modulation (HexMod) in human neuroimaing experiments. The strength of HexMod depends on the adaptation time constant, grid scale, movement speed, and length and straightness of behavioural trajectories. Accounting for these dependencies can improve experimental design and interpretation of non-invasive measure of grid cell activity across individuals.

## Method

### Experimental data overview

We analysed extracellular recordings of grid cells from six published rat datasets (Barry et al., 2012; Brun et al., 2008; Hafting et al., 2005; Ólafsdóttir et al., 2016; Sargolini et al., 2006; Vollan et al., 2025). Recordings were obtained from freely foraging rats in open-field arenas (square or circular), using tetrodes or silicon probes in the medial entorhinal cortex (MEC).

Only sessions lasting at least 10 minutes were included. All sessions were recorded during foraging from randomly scattered food in square open-field arenas (and a circular arena in Hafting et al., 2005). After filtering and grid cell identification procedure described in the following section, the dataset comprised 32 animals, 112 sessions, and 3,938 grid cells (Table 1). Of these, 1,171 (21 animals, 55 sessions) were conjunctive grid × direction cells (Rayleigh vector length above 0.3, see below). Figure 2 shows the conjunctive cells; all other analyses use all 3,938 cells. Due to multiple recording sessions in each animal, it is possible that the same cell could be recorded in more than one session and so counted more than once. We note that the interaction of the firing field locations and the trajectory taken in a given session is a critical factor in determining hexadirectional modulation (HexMod), which is counted uniquely.

**Table 1.** Summary of datasets analysed. Extracellular recordings of medial entorhinal cortex (MEC) grid cells from six published rat datasets. After filtering (sessions ≥ 10 min, square/circular arenas, gridness ≥ 0.3), the combined dataset included 32 animals, 112 sessions, and 3,938 grid cells.

| Dataset | #Animals | #Sessions | #Cells | Environment | Grid Scale |
| --- | --- | --- | --- | --- | --- |
| Brun et al. (2008) | 1 | 2 | 5 | Square, 220 × 220 cm (also 100 × 100 cm sessions) | ~60 – 150 cm |
| Barry et al. (2012) | 7 | 57 | 118 | Square, 100 × 100 cm | ~30 – 70 cm |
| Ólafsdóttir et al. (2016) | 6 | 24 | 61 | Square, ~100 × 100 cm | ~30 – 100 cm |
| Hafting et al. (2005) | 2 | 3 | 23 | Square 100 × 100 cm (also circular, diameter 100 & 200 cm) | ~30 – 70 cm |
| Sargolini et al. (2006) | 1 | 2 | 7 | Square, 100 × 100 cm (also 150 × 150 cm sessions) | ~70 – 120 cm |
| Vollan et al. (2025) | 15 | 24 | 3,724 | Square, 150 × 150 cm | ~50 – 150 cm |
| **Total** | **32** | **112** | **3,938** |  |  |

### Preprocessing of trajectories and spiking data

Position samples were resampled to 50Hz (*Δt* = 0.02s). Missing coordinates were linearly interpolated; trajectories were then smoothed with a 0.4s boxcar (moving-average) filter (20 frames at 50 Hz). Instantaneous velocity was computed from frame-to-frame differences; Speed 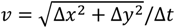 and movement direction θ = atan2(Δ*y*, Δ*x*) were derived and wrapped to [0,2*π*). Spike times were aligned to the 50Hz time base to form a per-sample spike array; spikes falling inside large gaps in tracking data (> 1.5 × nominal *Δt*) were excluded prior to binning. All analyses used non-stationary periods with speed ranges of 5 − 150*cm s*^−1^.

### Grid cells and conjunctive cell classification

For each cell, the firing rate map was calculated as the number of spikes divided by occupancy time in 3×3cm spatial bins, followed by Gaussian smoothing (σ=1bin). Spatial autocorrelograms were computed from the smoothed rate map and lightly smoothed (σ =1bin) to facilitate peak detection. Local maxima in the autocorrelogram were detected after masking the central peak, and an annular mask was drawn around six nearest peaks (inner radius=0.7, outer radius=1.3 of mean distance to peaks). The masked autocorrelogram was correlated with itself after rotations. Gridness was defined as the minimum of the correlation at 60° or 120° minus the maximum at 30°, 90° or 150°. Cells with gridness above 0.3 were considered as potential grid cells. Grid scale was defined as the mean distance of the six peaks from the centre. To determine the grid orientation of the cell, we measured the angle of each of six peaks relative to the x axis anticlockwise, then computed a circular mean of these angles relative to the 60° periodicity. By averaging orientation from multiple peaks, grid orientation could be more robustly estimated. We manually inspected the distribution of grid scales and orientation within each animal and clustered candidate grid cells into 1, 2 or 3 modules consisting of shared grid scale and orientation.

We then tested whether grid cells show additional directional tuning, i.e. conjunctive grid × head-direction cells. Directional firing rate was defined as the number of spikes divided by occupancy time in 5° movement direction bin. Of note, we used movement direction instead of head direction because movement direction tuning is required for HexMod. We smoothed the directional firing rate with a boxcar function (3 bins). The magnitude of Rayleigh vector length (R) indicates the strength of directional tuning, and the angle of the Rayleigh vector indicates the preferred direction:

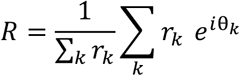

*r_k_* is the firing rate at k direction bin, and *θ_k_* is the associated direction. We defined conjunctive cell as having Rayleigh vector length greater than 0.3 following previous literature (Gardner et al., 2022). Tuning width is defined as the circular standard deviation of each cell’s directional firing rate map, 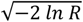. For comparison, a tuning curve proportional to 1 + *cos*(*θ* − *θ*_o_), with *θ*_o_ the preferred direction, has a circular standard deviation of 67.5°.

### Alignment and clustering of preferred directions

For each conjunctive cell, we classified the cell as aligned when its preferred direction lay within 15° of the nearest grid axis. Alignment with the grid axes was tested with a two-sided exact binomial test of the number of aligned cells against the 50% expected if preferred directions were unrelated to the grid.

For comparison with Doeller et al. (2010), we selected grid cells with their directional criterion: a Kullback-Leibler divergence of at least 0.15 (natural logarithm) between the directional firing rate map (Gaussian-smoothed, total SD about 10°, normalised to unit area) and a uniform distribution, using the earliest recording of each identified cell. This gave 890 cells from 21 animals, 888 of which were also conjunctive by our criterion. As in their Fig 1b, we computed the circular correlation coefficient of Fisher and Lee (1983) between grid orientation and preferred direction modulo 60°, both multiplied by six so that each spans the full circle. We tested whether it was greater than zero by randomly re-pairing preferred directions with grid orientations across cells (20,000 permutations; one-sided).

Clustering at 60° intervals around any angle was tested within each animal with a Rayleigh test on six times each cell’s preferred direction in room coordinates: multiplying by six makes directions 60° apart coincide, so that cells clustered around any six directions 60° apart share one angle. Because two recordings of the same cell share a preferred direction and would appear as clustering, this test counted each cell once. It was therefore restricted to the animals of Vollan et al. (2025), in which repeated recordings of a cell could be identified reliably from probe position, waveform and firing map; we used the earliest recording of each cell and the 12 animals with at least five conjunctive cells (1,082 cells). P values were Holm-corrected across the 12 animals, and the number of animals with p < 0.05 was compared with the 5% expected by chance (one-sided binomial test).

### Tuning width of conjunctive cells and hexadirectional modulation

To relate tuning width to six-fold modulation, we describe a directional tuning curve with a von Mises distribution, the circular equivalent of a Gaussian:

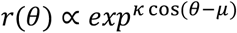

where *θ* is movement direction, *μ* is the preferred direction, and *κ* the concentration (larger *κ* means narrower tuning). Tuning width is expressed as the circular standard deviation 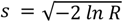, where *R* is the Rayleigh vector length of the tuning curve, the measure used to define conjunctive cells. For a von Mises curve *I*_1_(*κ*)/*I*_o_(*κ*) = *exp*(−*s*^2^/2), so⍰. A von Mises distribution can be expressed as a sum of n-fold angular components:

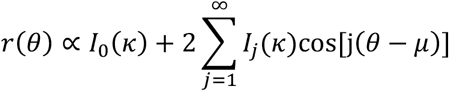

where *I_j_*(*κ*)is the modified Bessel function of the first kind of order j. The amplitude of the six-fold component cos[6(*θ* − *μ*)] relative to the mean is 2*I*_6_(*κ*)/*I*_o_(*κ*). For six cells with identical tuning whose preferred directions point along the six directions of the three grid axes (*μ*, *μ* + 60°, …, *μ* + 300°), the six-fold components of the six curves peak at the same directions and add up, whereas every component whose order is not a multiple of six cancels. The amplitude of the six-fold component relative to the mean remains the same as for a single cell, 2*I*_6_(*κ*)/*I*_o_(*κ*). The six-fold component falls steeply as tuning broadens (Supp Fig 5): for a von Mises curve it is 59% of the mean at a circular standard deviation of 15° (Fig 1C, narrow tuning), 0.3% at 43°, the width of the narrowest conjunctive cell, and 0.003% at 73°, the median width of the conjunctive cells.

**Supplementary Figure 5.**
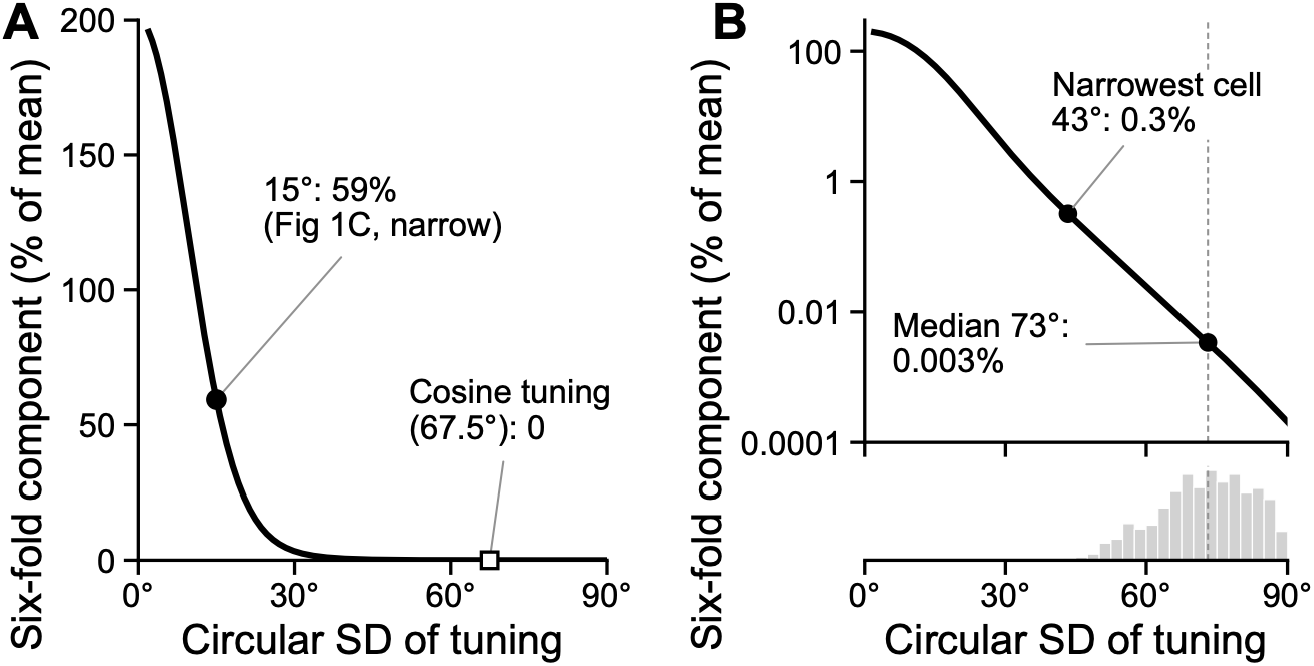
The six-fold component of a von Mises tuning curve falls steeply as tuning broadens. **A.** Six-fold component of a von Mises tuning curve (amplitude of the *cos*[6(*θ* − *μ*)] term as a percentage of the curve’s mean, 2I_6_(κ)/I_0_(κ); calculated, see Methods) against its circular standard deviation (SD). This is also the six-fold component of the summed activity of six identical cells, one per grid axis direction. Dot: 15°, the narrow tuning of Fig 1C (59%). Open square: cosine tuning, *r*(*θ*) ∝ 1 + *cos*(*θ* − *μ*), a different curve with a circular SD of 67.5° and no six-fold component (Fig 1C, broad tuning); a von Mises curve of the same width has 0.008%. **B.** The same on a logarithmic scale. Dots: 43°, the narrowest conjunctive cell (0.3%), and 73°, the median width of the 1,171 conjunctive cells (0.003%; dashed line). Below: number of conjunctive cells per 2.5° bin of circular SD, each computed from the Rayleigh vector length R of the cell’s movement-direction tuning as 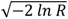. Because conjunctive cells were selected by R above 0.3, none can be broader than 89°.

Similarly, a cosine directional tuning curve, *r*(*θ*) ∝ 1 + *cos*(*θ* − *μ*) (circular standard deviation 67.5°), contains no six-fold component, so six grid-aligned cells with cosine tuning sum to a uniform rate (Fig 1C, broad tuning).

When cells with the same directional tuning shape and mean rate have imperfectly aligned preferred directions, the six-fold modulation of the summed population activity is reduced in proportion to their directional alignment. Specifically, it is given by 2*I*_6_(*κ*)/*I*_o_(*κ*) multiplied by the Rayleigh vector length of six times their preferred directions, |(1/*N*) *Σ_j_ exp*(6*iμ_j_*)|, where *N* is the number of cells and *μ_j_* the preferred direction of cell *j*.

### Grid field delineation

First, we smoothed the rate map using a Gaussian kernel with one-eighth of the grid scale to facilitate peak detection. Second, we detected local maxima and excluded the peaks with firing rate below baseline. Baseline firing rate was defined as the 40^th^ percentile of all bin values within the rate map. Third, a minimum inter-peak distance of two-thirds of the grid scale was enforced to prevent multiple detections within a single field. For each detected peak, the field radius was defined as the smaller of a) the distance at which at least 90% of bins within the radius exceeded baseline firing rate, or b) one-third of the grid scale.

### Quantifying firing rate adaptation: non-parametric model

To test whether a cell is less excitable after recent activity and identify the temporal profile of firing rate adaptation, we assessed the firing rate as a function of time elapsed since the previous field visit. We first identified the entry and exit of each grid field for given trajectory. The time elapsed since the exit of the previous field exit (*t_prev_*) was assigned to each time point (20 ms bin) until entry of the next field (Fig 3B top panel). *t_prev_* was clipped at the initial value inside the field.

To account for firing rate variability due to location and speed, we modelled the firing rate in each time bin (*i*) as follows:

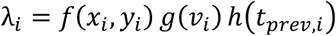

where *f*(*x_i_*, *y_i_*) is the spatial firing profile over 5 *cm* × 5 *cm* spatial bins, *g*(*v_i*) is the firing profile over eight quantile-based speed bins, and *h*(*t_prev_i_*) is the firing profile over four *t_prev_* bins [0,1), [1,2), [2,3), [3, ∞)*s*. Model parameters were estimated using an iterative fitting algorithm (Burgess et al., 2005) that alternately updated the spatial, speed, and temporal components to maximise the Poisson likelihood. Fitting continued for up to 100 iterations or until the increase in log-likelihood was negligible. At the end of fitting, *f*(*x,y*), *g*(*v*), *h*(*t_prev_*) were normalised so that the estimated firing rate for each factor alone matched the total numbers of spikes divided by the total duration. We then report the percent change of firing rate for each *t_prev_* relative to the mean firing rate of each cell. We ran 1-way ANOVA and post-hoc t-tests to test whether firing is lower for shorter *t_prev_* than long *t_prev_*.

### Quantifying firing rate adaptation: parametric model

One limitation of the non-parametric model in the previous section is that it assumes the same amount of adaptation whenever a grid field is visited. Adaptation-induced firing suppression should be stronger after a cell has received stronger spatial inputs, as would occur when the animal visits the centre of a field rather than its periphery. A second limitation of the model is the necessity for discretised binning of *t_prev_*. To tackle these two limitations, we developed a more realistic adaptation model which assumes the adaptation gain is an exponential time-decaying factor of *t_prev_*, exp(−*t_prev_*/τ) weighted by the summed location input from the previous field visit, w. Instantaneous location input was defined as a Gaussian function of distance between the animal’s position and the field centre, scaled by the mean firing rate of that field computed from the rate map (Fig 3B middle panel). The location-weighted time-decaying adaptation variable can be shown in Fig 3B bottom panel (adapt var).

We tested whether firing rate is explained by this adaptation variable, h(*t_prev_*) while controlling for speed and location using a Poisson multiplicative model:

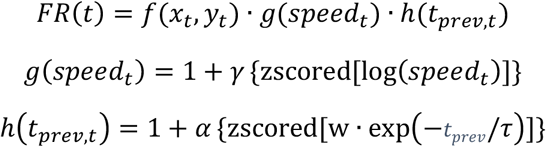

where *f*(*x_i_*, *y_i_*) is the spatial firing profile over 5*cm* × 5*cm* spatial bins. The influence of speed on firing rate is assumed to be logarithmic (Moghadam et al., 2025). We chose the time constant τ of 2 s in line with the slow adaptation time constant from in-vitro electrophysiology study of entorhinal layer II neurons (Alonso & Klink, 1993). We z-scored the speed and adaptation variable to improve stability of model fitting. We iteratively performed Newton’s methods to update *f*(*x_t_*, *y_t_*), *γ*, *α* to maximise Poisson log-likelihood, with bound condition that g(speed) and h(*t_prev_*) factor has to remain between 0 and 2. Positive *γ* indicates higher firing rate for faster speed. Negative *α* indicates lower firing rate for short *t_prev_*.

After fitting the model to entire trajectory data, we divided each recording into two halves (even and odd minutes) and fitted the model again using each half of data to check the robustness of adaptation effect. We assessed internal consistency by correlating fitted adaptation coefficients between even- and odd-minute subsets of each recording.

### Control analysis: prospective coding

If grid cells show prospective coding, i.e. encoding for location ahead of animals (Chaudhuri-Vayalambrone et al., 2023b; Csordas et al., 2024; De Almeida et al., 2012), the cell would start and stop firing slightly before entering/exiting the fields. Then a lower firing rate is expected for short *t_prev_*. To rule out this potentially confounding effect, we computed the rate map after shifting spike times relative to locations (−400 to 400 ms by 20 ms) and found the time shift which gives highest spatial information (Supp Fig 2A). We then fitted the adaptation model with shifted spikes.

### Quantifying firing rate adaptation across successive field visits

To test whether firing was reduced across successive grid-field visits, we performed a field-visit sequence analysis. Field visits were identified using the grid field delineation method described, then sorted by entry time for each cell. Candidate sequences consisted of four consecutive visits. We defined *T*_12_ as the time from exit of visit 1 to entry of visit 2, *T*_23_ as the time from exit of visit 2 to entry of visit 3, and *T*_34_ as the time from exit of visit 3 to entry of visit 4. Candidate sequences were retained when the following conditions were met. First, all three intervals satisfied 0 ≤ *T* ≤ 10 *s*, to ensure steady movements across fields with unbiased sampling of movement speed at visit 2 and 3. Second, sequences with the same field visited consecutively were excluded to avoid tortuous trajectories. Third, each field should be visited for at least 0.1s and there should be at least 2 expected spikes in field 2 and field 3 for robust estimation of firing rate ratio. For each cell, a baseline in-field firing rate was computed as the total number of spikes divided by total duration when an animal was visiting any of the grid fields. Expected spikes for each visit were computed as baseline in-field rate multiplied by visit duration.

For each candidate sequence, firing rates were computed for visits 2 and 3, and the firing rate ratio (FRR) was defined as

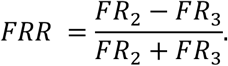

Positive values indicate lower firing during visit 3 than visit 2. Sequences were separated according to whether *T*_23_ < *T*_12_ or *T*_23_ > *T*_12_. For each cell, we computed the mean firing rate ratio in each timing condition. Cells contributed to the paired test only if they had valid sequences in both conditions.

### Relation between grid-aligned movements and time between fields

For firing rate adaptation to cause HexMod, grid-aligned movements should be associated with more firing rate adaptation (i.e. traversing two fields in quick succession) than grid-misaligned movements, which is not guaranteed for tortuous trajectory of freely moving animals. To assess the relationship between the instant movement direction and time between the fields, we classified each 20 ms time bin inside the field into the aligned (less than ±15° from nearest grid axis) and misaligned (more than ±15°). We then computed the mean temporal proximity as

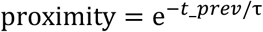

with τ = 2 s for all aligned and misaligned time bins. We used this non-linear transformed *t_prev_* instead of raw *t_prev_* to approximate the temporal profile of firing rate adaptation. We first compared the mean proximity difference between the aligned and misaligned movements using all cells. Then we divided grid cells into three groups of similar numbers of cells (1312-1313 cells/group) based on grid scale - small (< 62 cm), medium (62-86 cm), large (> 86 cm) and compared the mean proximity difference.

### Quantifying hexadirectional modulation

Following the method of Doeller et al. (2010), we defined a regressor for HexMod as *cos*[6(*θ*(*t*) − *φ*)], where *θ*(*t*) is the instantaneous running direction at time t, and *ϕ* is the grid orientation. In human studies, fMRI activity is simply regressed with this six-fold regressor. However, a single grid cell’s firing is strongly modulated by location and speed, and a simple regression method cannot account for unbalanced sampling of location and speed by running direction in foraging animals. For example, animals can approach some locations more frequently from a particular direction and the interaction between location and direction can cause either false positive or false negative HexMod. Similarly, if animals move faster along the boundary of the square environment, higher firing rate at cardinal directions due to speed can obscure HexMod.

To overcome these limitations, we developed a multiplicative Poisson model tailored to spike count data (Burgess et al., 2005). By modelling firing rates as products of independent factors – position, speed, and six-fold directional modulation. In the log domain, the formulation is equivalent to a generalised linear model with additive predictors, closely related to prior models of MEC firing (Hardcastle et al., 2017). The multiplicative Poisson model therefore provides a principled extension of cosine regression, yielding more reliable estimates of HexMod for individual neurons under naturalistic behaviour.

Instantaneous firing rate was modelled as a multiplication of the location factor (5 cm bin), z-scored logarithm of speed, and six-fold regressor.

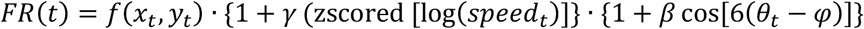

We iteratively performed Newton’s methods to update *f*(*x_t_*, *y_t_*), *γ*, *β* to maximise Poisson log-likelihood, with bound condition that speed and six-fold factor should remain between 0 and 2. Negative β values reflect suppressed firing during aligned movement, as the adaptation hypothesis predicts; positive values reflect stronger firing during aligned movement.

### Simulation: overview

We simulated the grid cell population activity for human spatial memory and rodent free foraging experiments, and applied the commonly used cross-validated hexadirectional analysis to examine the detectability of grid-like activity in the typical sample size (30 participants). For simplicity, we modelled a macroscopic signal (e.g. fMRI signal in a single voxel) as the sum of firing rate of multiple grid cells (n=10^4) which shares the same grid orientation and scale. Grid scale was fixed as either 40% or 70% of environmental size - typical smallest and largest grid scales found in the rodent experiment.

### Simulation: number of grid cells

Total number of neurons in the human entorhinal cortex is estimated as 7*10^6 (Gómez-Isla et al., 1996; Insausti & Amaral, 2012) and in rodents, approximately 20% of cells in superficial layers of medial entorhinal cortex showed significant gridness, including the conjunctive grid cells (Diehl et al., 2017). One in-vivo study in humans found 12 out of 89 (13%) recorded entorhinal units showed grid-like property (Jacobs et al., 2013). Human entorhinal cortex has a total volume of 1.1*10^3 mm^3^ (Oltmer et al., 2022) and thickness of 2.5~3 mm (Feczko et al., 2009), resulting in ~70 voxels of 2.5 mm isotopic voxel size. Assuming 10% of whole entorhinal neurons are grid cells, 10^4 grid cells are expected per voxel and this number is used for simulation.

### Simulation: trajectories

We created realistic trajectories for an object-location memory paradigm that is commonly used in human experiments. For each 10 min experimental block (sampling rate = 10 Hz), we selected multiple fixed target locations in a square environment with a length of 100 virtual metres (vm) and choose the random start locations where participants could move to the proximity of each target location from a wide range of movement direction (360°) and distance (30~70 vm). We used a fixed speed (16 vm/s) that is comparable to rat data (rat’s mean movement speed was 19 cm/s in 100 cm environments) and published human experiments; c.f. (Convertino et al., 2023) used 20 vm/s for an environment size of 100 vm). We also added intertrial intervals (2 s), cue (1 s) and rotation (0.8~3.2 s) periods similar to human experiments, but only movement periods were used for analysis. We created four blocks of trajectories for each of 30 virtual participants.

### Simulation: firing rate of individual grid cells

Grid cells (GC) firing was modelled as a multiplication of the location-dependent baseline rate map (see details in the next section) and the adaptation factor based on time between the field as follows:

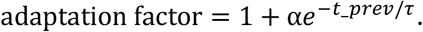

Adaptation strength α was set as 0.1, which results in ~8% reduction in firing rate for short intervals (0.5s) compared to the longest intervals, based on percent firing change observed in real grid cells from a non-parametric model of adaptation (Fig 3C). Time constant τ was set as 2 s. We didn’t add a speed modulation because we used a trajectory with constant speed.

### Baseline rate map of simulated grid cells

Grid cells have been previously modelled as a sum of three plane waves 60° apart, which places the centres of each grid field at the vertex of perfect triangular lattice with homogeneous firing rate for each field. However, individual grid fields show large and consistent variability in firing rates (Ismakov et al., 2017) and local distortion in spatial arrangement of grid fields are observed in various studies dependent on environmental boundaries (Duncan et al., 2025; Krupic et al., 2015; Ying et al., 2023), reward (Boccara et al., 2019) and experience (Duncan et al., 2025), see example grid cells in Supp Fig 3A. Inhomogeneous firing rates and spatial deviations from perfect lattices would reduce HexMod, therefore we quantified these two characteristics from real grid cells and applied it when simulating the baseline location firing of synthetic grid cells. We separately estimated the spatial deviation and firing rate inhomogeneity parameters for small (grid scale < 40% of environment size) and large (grid scale >60% of environment size) grid cells due to the different numbers of fields.

Variability in the mean firing rate of each grid field was characterised at two levels, across-cell and within-cell. Spatial deviation of each field was quantified as the distance between the centre of delineated fields and the best-fitting triangular lattice (white lines in Supp Fig 3A). We fitted a gamma distribution to spatial deviation, across- and within-cell firing rate distributions from real grid cells and estimated the shape and scale parameters (Supp Fig 3B,C,D). Based on gamma fit parameters, we added random jittering in spatial location and field firing rate to perfect hexagonal rate maps, resulting in realistic rate maps. Random jittering in field locations created some cells with low gridness, and we only retained the cells with gridness higher than 0.3. Shape and scale parameters for spatial jitter and firing rate inhomogeneity, example grid cells’ firing rate maps can be found in Supp Fig 3E.

### Measuring 6-fold and control symmetry in simulated data

Hexadirectional modulation of grid cell populations is modelled as cosi6(*θ* − *ϕ*)*j*, where *θ* is movement direction and *ϕ* is a grid orientation. We estimated the grid orientation from training data (three out of four blocks) using the quadrature filter approach which uses the ratio of regression coefficients for cos(6*θ*) and sin(6*θ*) regressors (Doeller et al., 2010). Stationary periods (speed <=5vm/s) were excluded from analysis. We then computed the regression coefficient for cosi6(*θ* − *ϕ*)*j* in the remaining data (1 block). This procedure is repeated for each unique run, and the averaged regression coefficient is defined as the strength of HexMod in that participant. This single value of HexMod strength is tested against 0 in a group of participants (one-sided t-test). For control N-fold symmetry analysis, we repeated the procedure with cosi*N*(*θ* − *ϕ*)*j*, where N is from 1 to 7.

For 6-fold symmetry, we also measured the mean activity value for 12 direction bins centred at 0, 30, 60, … 330° relative to the true grid orientation *ϕ* which is known for simulated data.

To account for jittering in GC rate maps and random selection of movement trajectory in each experiment, we repeated the simulation 50 times with different random seeds and counted how many simulation resulted in significant N-fold symmetry at the group level (p<0.05).

### Measuring HexMod strength while controlling for location

If all grid cells have the perfect homogeneous hexagonal rate map with evenly distributed grid offsets, firing rate variability due to location is expected to be negligible when cells are summed together. However, due to slight deviation from the perfect lattice of actual/simulated of each grid cell, the summed rate map shows location-dependent firing rate which can confound the estimate of firing rate for each movement direction. To mitigate this issue, we estimated the location-controlled directional firing by fitting a multiplicative model:

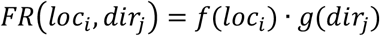

where *f*(*loc_i_*) is the spatial firing over 5 *cm* × 5 *cm* spatial bins and *g*(*dir_j_*) is the directional firing at each binned direction. We iteratively updated *f*(*loc_i_*) and *g*(*dir_j_*) by minimising the error,

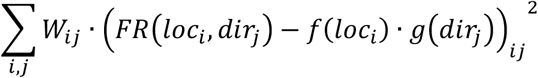

*W_ij_* is the occupancy of i-th location and j-th directional bin.

To estimate the firing rate for aligned and misaligned movement directions relative to the ground truth grid orientation, we used 30° bins. For quadrature filter approach in which grid orientation has to be estimated from a part of data, we used 5° bins in each training and testing data.

### Simulation: speed-dependency of HexMod strength

Movement speed influences the time interval between the grid field visits, and hence the adaptation gain. To measure the effect of speed alone, we simulated a simplified case with ideal hexagonal grid fields (homogeneous firing rate and perfect hexagonal geometry without any jitter). We generated one small module (scale: 30 vm,) and one large module (scale: 70 vm) each with 1024 grid cells. We used a continuous long straight run from one wall to another wall in a square environment (size: 100 vm). Movement direction changed randomly at each wall. We applied a constant speed for each simulation, ranging from 5 vm/s to 150 vm/s with 4 vm/s intervals. Each trajectory length was four blocks of 10 minutes. We measured the mean population activity during aligned and misaligned movement relative to the true grid orientation. The difference between the aligned and misaligned condition was divided by mean activity and then multiply by 100, resulting in a percent signal change.

### Optimal speed that maximises HexMod

The reduction in firing rate is expected to be proportional to proximity, defined as exp(−*t_prev_*/τ), where *t_prev_* is time between grid fields and τ is a time constant. Time between the fields in aligned direction is approximately 1*grid scale divided by speed v, and the time for misaligned direction is 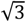 times of that.

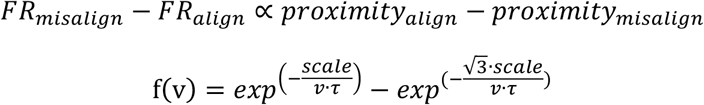

Let

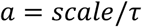

Then

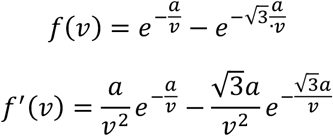

Set *f*′(*v*) = 0:

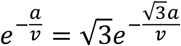

Taking logs,

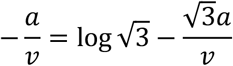

Therefore

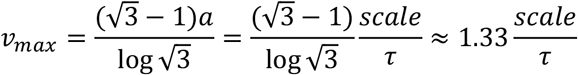

## Data and code availability

### Data

This paper analyses existing, publicly available datasets. The recordings of Vollan et al. (2025) are available on EBRAINS (doi:10.25493/R5FR-EDG; licence CC BY-SA 4.0). The recordings of Hafting et al. (2005), Sargolini et al. (2006) and Brun et al. (2008) are available in the NIRD Research Data Archive (doi:10.11582/2017.00017, doi:10.11582/2017.00019 and doi:10.11582/2018.00015, respectively; licence CC BY 4.0). The sessions of Barry et al. (2012) and Ólafsdóttir et al. (2016) analysed here are available on Zenodo (doi:10.5281/zenodo.23477461; licence CC BY 4.0).

### Code

All original code is available at https://github.com/gullirg/hexmod-grid-cell-adaptation (MIT licence). It downloads the six datasets and reproduces every figure and statistic reported in this paper, including the simulations.

## Acknowledgements

Funding support for this work was received from a Wellcome Principal Research Fellowship (222457/Z/21/Z) (NB), a BBSRC Project Grant (UKRI1950) (NB, CB), a BBSRC Research Grant (BB/Y011775/1) (CB), a UKRI Frontier Research grant (EP/X023060/1) (DB) and a UCL Division of Psychology and Language Sciences Executive Training Demonstrator Studentship (GR).

## Author Contributions

GR, MK, and NB designed the study, with input from DB, CB. GR and MK analysed the data and wrote the original draft. CB provided data. ZJ, DB provided inputs to analyses. All authors edited the manuscript.

## Declaration of generative AI and AI-assisted technologies in the manuscript preparation process

During the preparation of this work, the authors used ChatGPT (OpenAI), Gemini (Google) and Claude (Anthropic) for aiding coding and editing text. The authors reviewed and edited the output as needed and take full responsibility for the content of the published article.

